# *distinct*: a novel approach to differential distribution analyses

**DOI:** 10.1101/2020.11.24.394213

**Authors:** Simone Tiberi, Helena L Crowell, Pantelis Samartsidis, Lukas M Weber, Mark D Robinson

## Abstract

We present *distinct*, a general method for differential analysis of full distributions that is well suited to applications on single-cell data, such as single-cell RNA sequencing and high-dimensional flow or mass cytometry data. High-throughput single-cell data reveal an unprecedented view of cell identity and allow complex variations between conditions to be discovered; nonetheless, most methods for differential expression target differences in the mean and struggle to identify changes where the mean is only marginally affected. *distinct* is based on a hierarchical non-parametric permutation approach and, by comparing empirical cumulative distribution functions, identifies both differential patterns involving changes in the mean, as well as more subtle variations that do not involve the mean. We performed extensive bench-marks across both simulated and experimental datasets from single-cell RNA sequencing and mass cytometry data, where *distinct* shows favourable performance, identifies more differential patterns than competitors, and displays good control of false positive and false discovery rates. *distinct* is available as a Bioconductor R package.

## Background

Technology developments in the last decade have led to an explosion of high-throughput single-cell data, such as single-cell RNA sequencing (scRNA-seq) and high-dimensional flow or mass cytometry data, allowing researchers to investigate biological mechanisms at single-cell resolution. Single-cell data have also extended the canonical definition of differential expression by displaying cell-type specific responses across conditions, known as differential state (DS) [31], where genes or proteins vary in specific sub-populations of cells (e.g., a cytokine response in myeloid cells but not in other leukocytes [12]). Classical bulk differential expression methods have been shown to perform well when used on single-cell measurements [24, 25, 30] and on aggregated data (i.e., averages or sums across cells), also referred to as pseudo-bulk (PB) [6, 31]. However, most bulk and PB tools focus on shifts in the means, and may conceal information about cell-to-cell heterogeneity. Indeed, single-cell data can show more complex variations (Figure 1 and Supplementary Figure 1); such patterns can arise due to increased stochasticity and heterogeneity, for example owing to oscillatory and unsynchronized gene expression between cells, or when some cells respond differently to a treatment than others [14, 30]. In addition to bulk and PB tools, other methods were specifically proposed to perform differential analyses on single-cell data (notably: *scDD* [14], *SCDE* [13], *MAST* [10], *BASiCS* [9, 28, 29] and mixed models [26]). Nevertheless, they all present significant limitations: BASiCS does not perform cell-type specific differential testing between conditions, scDD does not directly handle covariates and biological replicates, while PB, SCDE, MAST and mixed models performed poorly in previous benchmarks when detecting differential patterns that do not involve the mean [6, 14].

**Figure 1:**
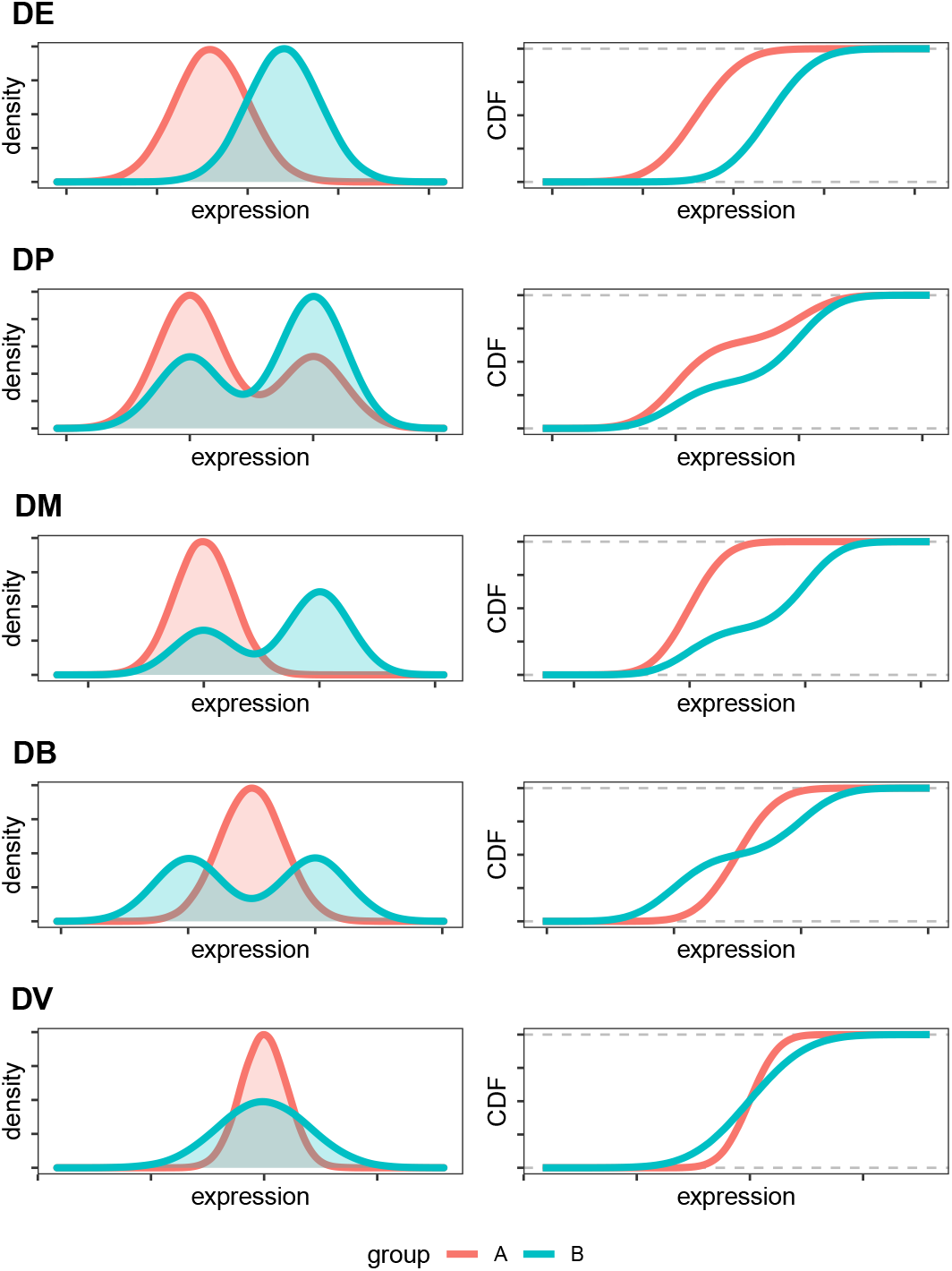
Cumulative distribution functions (CDFs) unravel differences between distributions. Density (left panels) and CDF (right panels) of five differential patterns: differential variability (DV), and the four proposed by Korthauer et. al. [14]: differential expression (DE), differential proportion (DP), differential modality (DM), and both differential modality and different component means (DB).

## Results

### *distinct* ‘s full distribution approach

To overcome these challenges, we developed *distinct*, a flexible and general statistical methodology to perform differential analyses between groups of distributions. *distinct* is particularly suitable to compare groups of samples (i.e., biological replicates) on single-cell data.

Our approach computes the empirical cumulative distribution function (ECDF) from the individual (e.g., single-cell) measurements of each sample, and compares ECDFs to identify changes between full distributions, even when the mean is unchanged or marginally involved (Figure 1 and Supplementary Figure 1). First, we compute the ECDF of each individual sample; then, we build a fine grid and, at each cut-off, we average the ECDFs within each group, and compute the absolute difference between such averages. A test statistic, *s*^*obs*^, is obtained by adding these absolute differences.

More formally, assume we are interested in comparing two groups, that we call *A* and *B*, for which *N*_*A*_ and *N*_*B*_ samples are available, respectively. The ECDF for the *i*-th sample in the *j*-th group, is denoted by 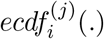, for *j* ∈ {*A, B*} and *i* = 1, …, *N*_*j*_. We then define *K* equally spaced cut-offs between the minimum, *min*, and maximum, *max*, values observed across all samples: *b*_1_, …, *b*_*K*_, where *b*_*k*_ = *min* + *k × l*, for *k* = 1, …, *K*, with *l* = (*max − min*)*/*(*K* + 1) being the distance between two consecutive cut-offs. We exclude *min* and *max* from the cut-offs because, trivially, 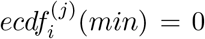 and 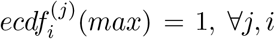. At every cut-off, we compute the absolute difference between the mean ECDF in the two groups; our test statistic, *s*^*obs*^, is obtained by adding these differences across all cut-offs:

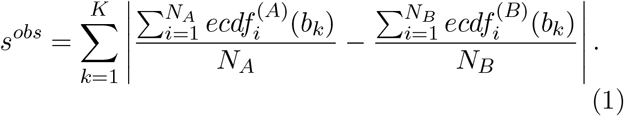

Note that in differential state analyses, these operations are repeated for every gene-cluster combination.

Intuitively, *s*^*obs*^, which ranges in [0, ∞), approximates the area between the average ECDFs, and represents a measure of distance between two groups of densities: the bigger *s*^*obs*^, the greater the distance between groups. The number of cut-offs *K*, which can be defined by users, is set to 25 by default, because no detectable difference in performance was observed when further increasing it (data not shown). Note that, although at each cut-off we compute the average across each group’s curves, ECDFs are computed separately for each individual sample, therefore our approach still accounts for the within-group variability; indeed, at a given thresh-old, the average of the sample-specific ECDFs differs from the group-level ECDF (i.e., the curve based on all individual measurements from the group). The null distribution of *s*^*obs*^ is then estimated via a hierarchical non-parametric permutation approach (see Methods). A major disadvantage of permutation tests, which often restricts its usage on biological data, is that too few permutations are available from small samples. We overcome this by permuting cells, which is still possible in small samples, because there are many more cells than samples. In principle, this may lead to an inflation of false positives due to lack of exchangability (see Methods); nonetheless, in our analyses, *distinct* provides good control of both false positive and false discovery rates.

Importantly, *distinct* is general and flexible: it targets complex changes between groups, explicitly models biological replicates within a hierarchical framework, does not rely on asymptotic theory, avoids parametric assumptions, and can be applied to arbitrary types of data. Additionally, *distinct* can also adjust for sample-level cell-cluster specific covariates (i.e., whose effect varies across cell clusters), such as batch effects. In particular, *distinct* fits a linear mixed effects model with the input data (e.g., normalized counts) as response variable, nuisance covariates as fixed effects, and samples as random effects. The method then removes the estimated impact of fixed effect covariates, and performs differential testing on these normalized values (see Methods).

Furthermore, to enhance the interpretability of differential results, *distinct* provides functionalities to compute (log) fold changes between conditions, and to plot densities and ECDFs, both for individual samples and at the group-level.

Note that, although *distinct* and the Kolmogorov-Smirnov [17] (KS) test share similarities (they both compare distributions via non-parametric tests), the two approaches present several conceptual differences. Firstly, the KS considers the maximum distance between two ECDFs, while our approach estimates the overall distance between ECDFs, which in our view is a more appropriate way to measure the difference between distributions. Secondly, the KS test only compares two individual densities, while our framework compares groups of distributions. Thirdly, while the KS statistic relies on asymptotic theory, our framework uses a permutation test. Finally, a comparison between *distinct* and *scDD* [14] based on the KS test (labelled *scDD-KS*) shows that our method, compared to the KS test, has greater statistical power to detect differential effects and leads to fewer false discoveries (see Simulation studies).

### Simulation studies

We conducted an extensive benchmark, based on scRNA-seq and mass cytometry simulated and experimental datasets to investigate *distinct* ‘s ability to identify differential patterns in sub-populations of cells.

First, we simulated droplet scRNA-seq data via *muscat* [6] (see Methods). We ran five simulation replicates for each of the differential profiles in Figure 1, with 10% of the genes being differential in each cluster, where DE (differential expression) indicates a shift in the entire distribution, DP (differential proportion) implies two mixture distributions with different proportions of the two components, DM (differential modality) assumes a unimodal and a bimodal distribution, DB (both differential modality and different component means) compares a unimodal and a bimodal distribution with the same overall mean, and DV (differential variability) refers to two unimodal distributions with the same mean but different variance (Figure 1 and Supplementary Figure 1). Each individual simulation consists of 4,000 genes, 3,600 cells, separated into 3 clusters, and two groups of 3 samples each, corresponding to an average of 200 cells per sample in each cluster.

We considered five different normalization approaches: counts per million (CPMs), *scater* ‘s logcounts [18], *linnorm* [33], *BASiCS* [9, 28, 29], and residuals from variance stabilizing normalization from *sctransform* (vstresiduals) [11]. We compared *distinct* to several PB approaches from *muscat*, based on *edgeR* [23], *limma-voom* and *limma-trend* [22], which emerged among the best performing methods for differential analyses from scRNA-seq data [6, 25]. We further considered three methods from *muscat* based on mixed models (MM), namely *MM-dream2, MM-vstresiduals* and *MM-nbinom* (see Methods). Finally, we included *scDD* [14], which is conceptually similar to our approach: *scDD* implements a non-parametric method to detect changes between individual distributions from scRNA-seq, based on the Kolmogorov-Smirnov test, *scDD-KS*, and on a permutation approach, *scDD-perm*. For *scDD-perm* we used 100 permutations to reduce the computational burden.

In all scenarios and on all five input data, *distinct* shows favourable performance: it has good statistical power while controlling for the false discovery rate (FDR) (Figure 2). In particular, for DE, DP and DM, *distinct* has similar performance to the best performing competitors (*edgeR*.*linnorm* and *limma-trend*.*logcounts*), while for DB and DV, it achieves significantly higher true positive rate (TPR), especially when using *log-counts*. PB methods in general perform well for differential patterns involving changes in the mean (DE, DP and DM), but struggle to identify DB and DV patterns. *scDD* provides good TPR across all patterns when using the KS test on vstresiduals (*scDD-KS*.*vstresiduals*), while the TPR is significantly reduced when using other inputs and with the permutation approach(*scDD-perm*); however, *scDD* methods (in particular, *scDD-KS*.*vstresiduals*) also show a significant inflation of the FDR. In contrast, MM methods provide good control of the FDR but have low statistical power in all differential scenarios. We also investigated how normalization influences each method’s results (Supplementary Figure 2): *distinct* appears to be the least affected method and displays the smallest variation across normalization inputs, possibly due to its non-parametric structure, which can more flexibly accommodate various inputs.

**Figure 2:**
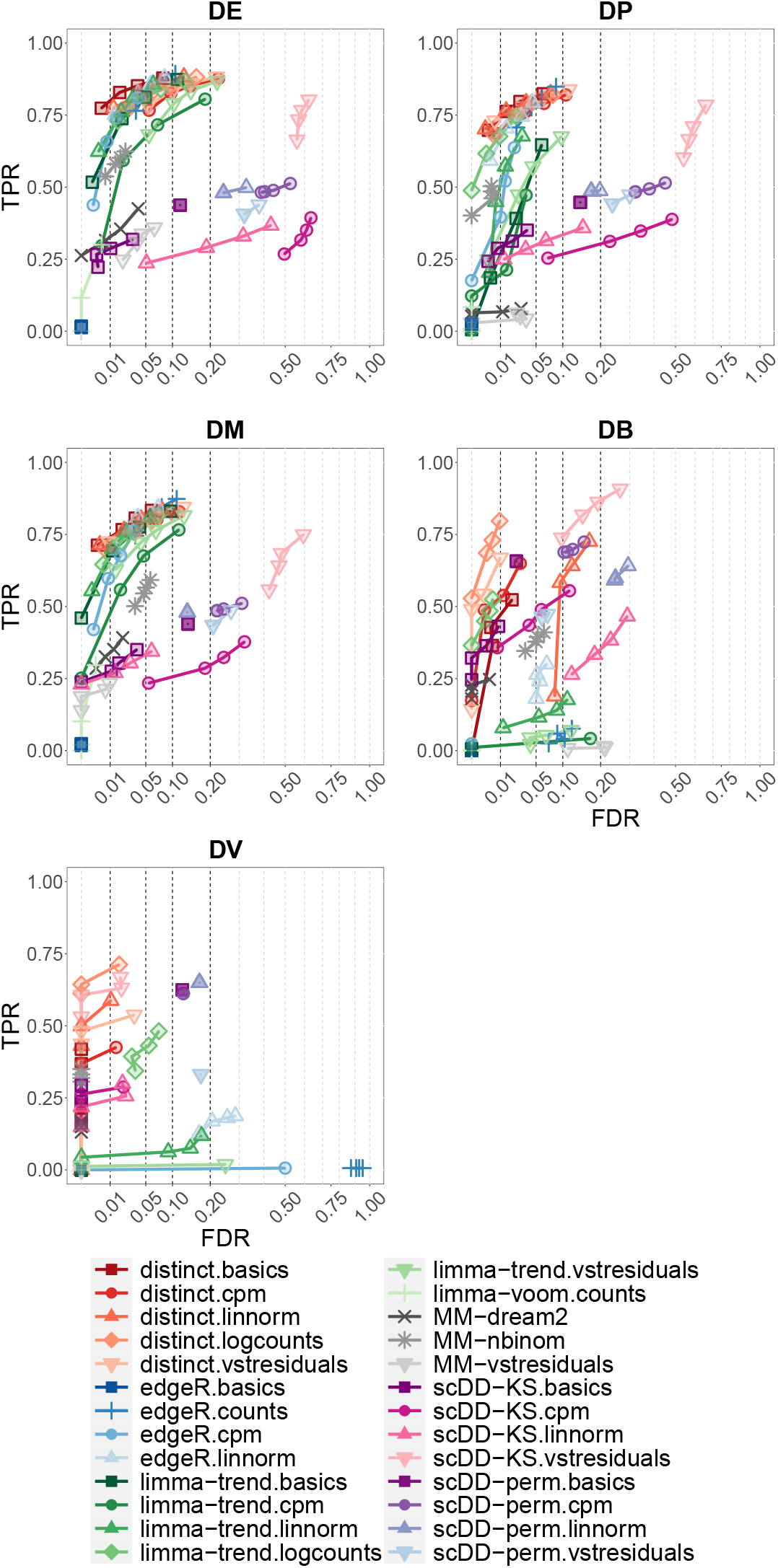
*distinct* identifies various differential patterns and controls for the FDR. TPR *vs*. FDR in *muscat* simulated data; DE, DP, DM, DB and DV refer to the differential profiles illustrated in Figure 1. Circles indicate observed FDR for 0.01, 0.05, 0.1 and 0.2 significance thresholds. Results are averages across the five simulation replicates. Each individual replicate consists of 4,000 genes, 3,600 cells, separated into 3 clusters, and two groups of 3 samples each, corresponding to an average of 200 cells per sample in each cluster.

We further simulated five null simulation replicates with no differential patterns; again with each simulation having 4,000 genes, 3,600 cells, 3 cell clusters and two groups of 3 samples each. In the null simulated data, only *limma-trend*.*basics* and *limma-trend*.*cpm* present a mild inflation of false positives, while MM and, particularly, *edgeR*.*basics* lead to overly conservative p-values; instead, *distinct* and *scDD* show approximately uniform p-values for all types of input data (Figure 3).

**Figure 3:**
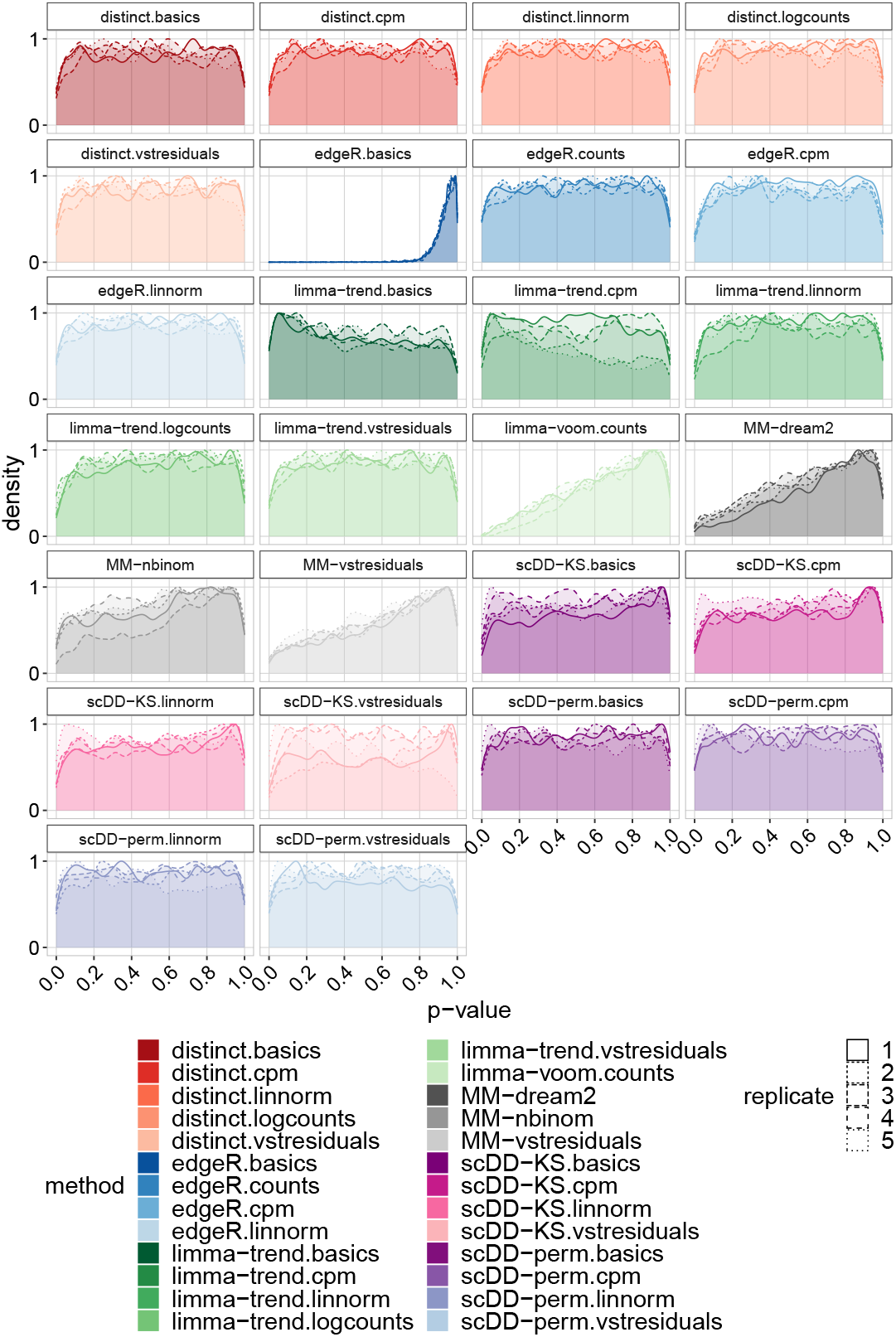
*distinct* has uniform null p-values. Density of raw p-values in *muscat* null simulated data; each replicate represents a different null simulation. Each individual replicate consists of 4,000 genes, 3,600 cells, separated into 3 clusters, and two groups of 3 samples each, corresponding to an average of 200 cells per sample in each cluster.

We also extended previous simulations to add a cell-type specific batch effect (i.e., a batch effect that affects differently each cell-type) [6, 16]. In particular, we simulated 2 batches, that we call *b*_1_ and *b*_2_, with one group of samples having two samples associated to *b*_1_ and one to *b*_2_, and the other group of samples having two samples from batch *b*_2_ and one from *b*_1_. Differential results are substantially unchanged (Supplementary Figure 3), which shows *distinct* can effectively remove nuisance confounders.

Furthermore, we performed various sensitivity analyses and investigated how results are affected when varying: i) the number of cells, ii) the library size, iii) the dispersion parameter, iv) the fraction of significant genes, and v) the sample sizes in each group. In particular, we simulated 50, 100, 200 (as in the original simulation) and 400 cells per sample in each cluster. We further modified the library size and dispersion parameters of the negative binomial model used by *muscat* to simulate scRNA-seq data, influencing the mean expression and cell-to-cell variability respectively, by considering values 1/5, 1/2, 2 and 5 times as big as those used in the original simulation. In addition, we varied the percentage of simulated differential genes as 1, 5, 10 (as in the original simulation) and 20%, and considered various unbalanced designs by comparing two groups of different sample sizes: 3 *vs*. 2, 4 *vs*. 3, and 5 *vs*. 3. Overall, increasing the number of cells or the library size and decreasing the dispersion have a positive impact on the performance of all methods, by improving their ability to detect differential effects (i.e., true positive rate); nonetheless, none of these factors seem to affect the relative ranking of methods, which remains globally stable (Figure 4 and Supplementary Figures 4-5). In addition, changing the fraction of significant genes and considering unbalanced designs does not appear to introduce systematic changes in performance (Supplementary Figures 6-7). Note that, in these sensitivity analyses, we excluded MM models due to the high computational cost and low statistical power displayed in the previous analyses.

**Figure 4:**
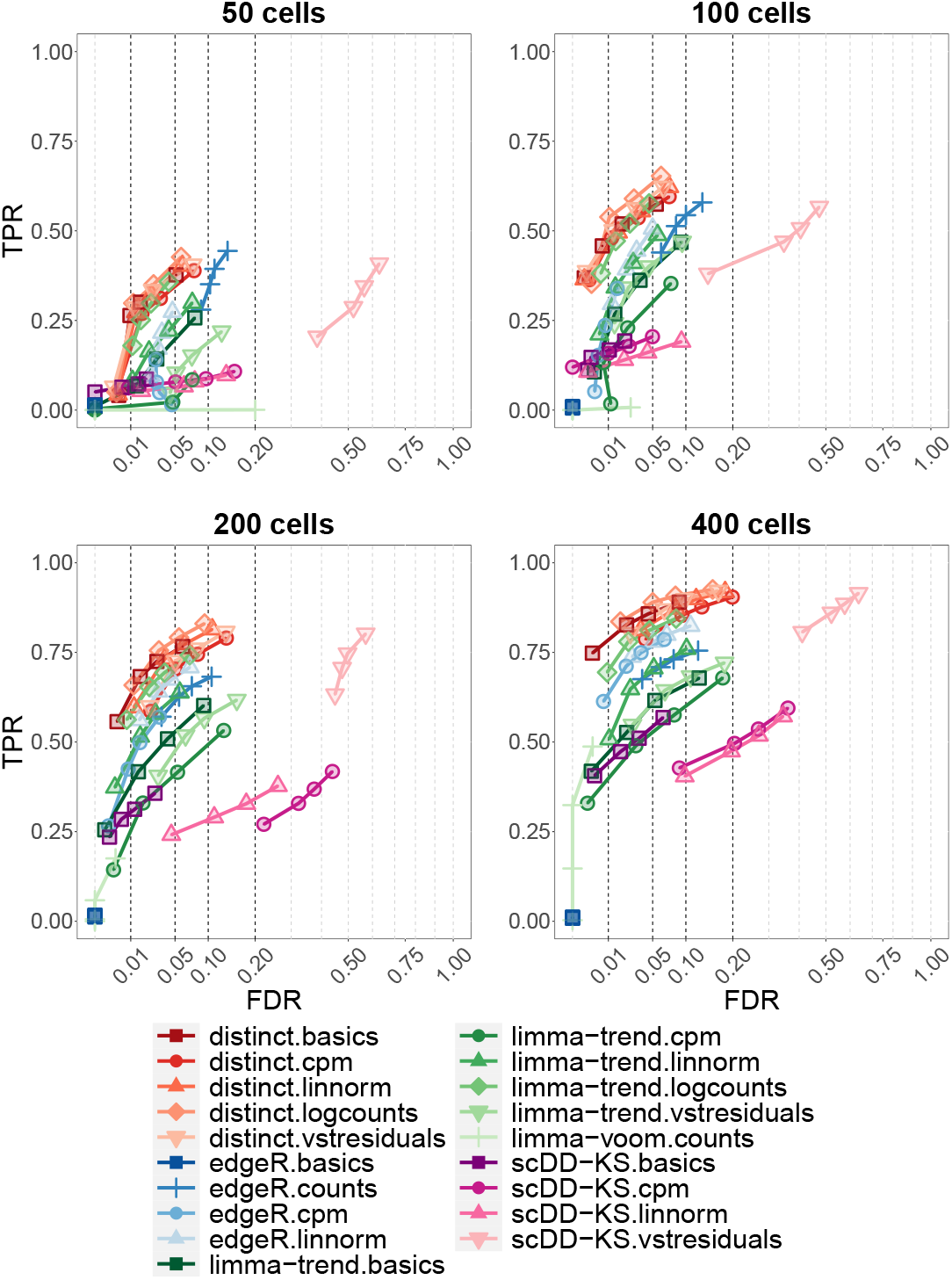
*distinct* achieves good performance when varying the number of available cells. TPR *vs*. FDR in *muscat* simulated data; with 50, 100, 200 and 400 cells per cluster-sample combination, corresponding to a total of 900, 1,800, 3,600 and 7,200 cells, respectively. Results are aggregated over the five replicate simulations of each differential type (DE, DP, DM, DB and DV), contributing in equal fraction. Each individual simulation replicate consists of 4,000 genes, 3 cell clusters and two groups of 3 samples each. Circles indicate observed FDR for 0.01, 0.05, 0.1 and 0.2 significance thresholds. Note that *scDD-perm* and MM were excluded from this analysis due to their computational cost.

From a computational perspective, *distinct* required an average time of 3.2 to 4.5 minutes per simulation, which is higher than PB methods (0.1 to 0.2 minutes) and *scDD-KS* (0.5 to 0.7 minutes), but significantly lower than MM approaches (29.4 to 297.3 minutes) and *scDD-perm* (544.7 to 2085.6 minutes) (Figure 5 and Supplementary Table 1). All methods were run on 3 cores, except PB approaches, which used a single core, because they do not allow for parellel computing.

**Table 1:** Percentage of unique gene/cell-type identifications that are unique to each method. Since methods return significantly different number of significant results, for each method, we selected the most significant 1,000 results. For every method, we then compute the fraction of such results that are unique, i.e., not in common with the top 1,000 results returned by any other method.

| method | % of unique results |
| --- | --- |
| distinct.logcounts | 0.3 |
| distinct.basics | 0.8 |
| limma-trend.logcounts | 0.9 |
| distinct.cpm | 1.0 |
| distinct.vstresiduals | 1.1 |
| edgeR.linnorm | 1.2 |
| limma-trend.vstresiduals | 1.5 |
| limma-trend.basics | 1.5 |
| edgeR.counts | 1.7 |
| edgeR.basics | 3.0 |
| distinct.linnorm | 3.6 |
| limma-trend.linnorm | 3.7 |
| limma-voom.counts | 5.6 |
| edgeR.cpm | 10.4 |
| limma-trend.cpm | 26.8 |

**Figure 5:**
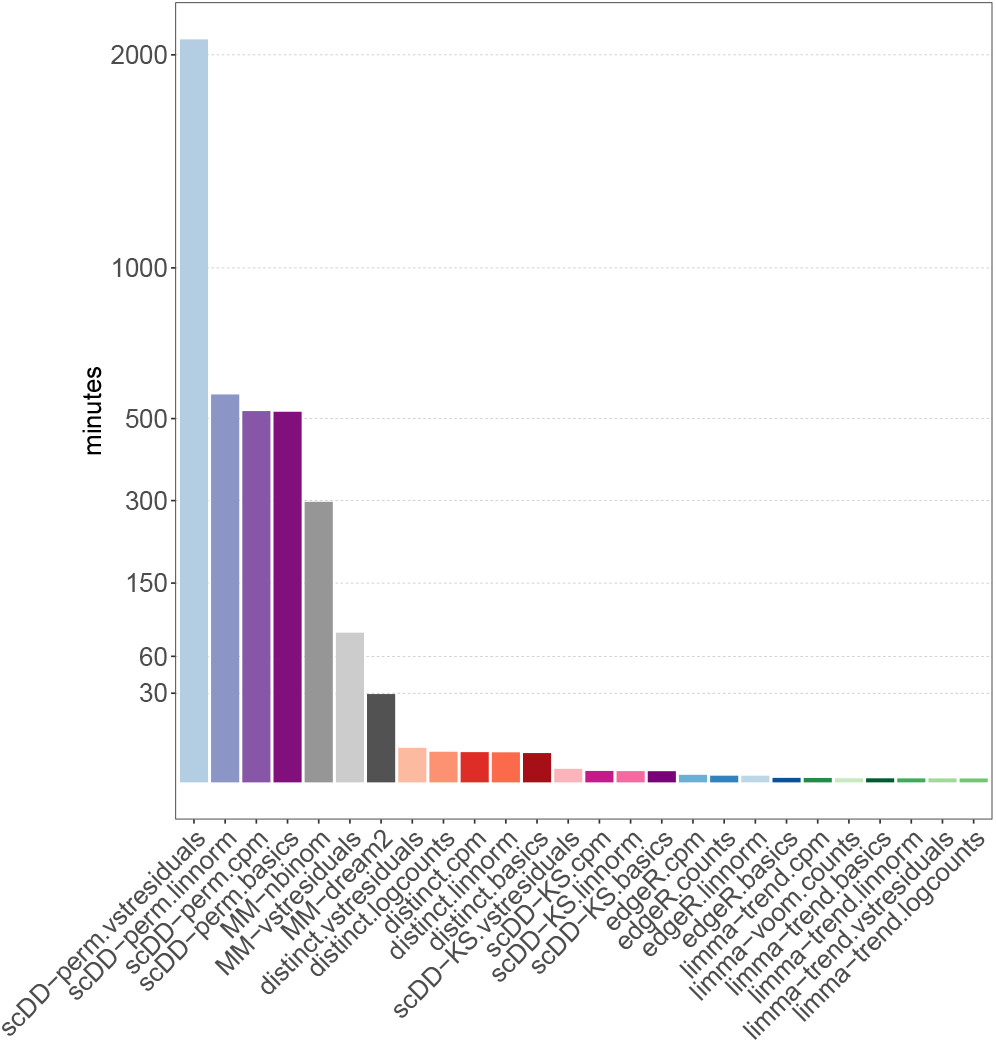
*distinct* requires more computational resources than PB and *scDD-KS* methods, but significantly less than MM and *scDD-perm* models. Average computing time, expressed in minutes, in *muscat* main simulations (Figures 2-3). For each method, times are averaged across simulation types (DE, DP, DM, DB, DV and null) and, for each type, across the five replicate simulations; in each replicate 3,600 cells are available (200, on average, per cluster-sample combination). *distinct*, MM and *scDD* models were run on 3 cores, while pseudo-bulk methods based on *edgeR* and *limma* used a single core because they do not allow for parellel computing. Note that *scDD-perm* requires much longer on vstresiduals than on the other normalized data, because *scDD* performs differential testing on non-zero values: vstresiduals, (unlike linnorm, cpm and basics normalized data) are not zero-inflated and, therefore, many more cells have to be used for differential testing.

We also considered an alternative popular droplet scRNA-seq data simulator, *SplatPOP* [2], which represents a generalization of *Splatter* [34], that allows multi-sample multi-group synthetic data to be generated. In particular, we simulated 20,345 genes from a human genome with two groups of 4 samples each, and 100 cells per sample, belonging to the same cluster of cells, for a total of 800 cells across all samples. We ran 8 differential simulations, with 10% of genes truly differential between groups, by varying the location (*de*.*facLoc*) and scale (*de*.*facScale*) differential parameters, mainly affecting the mean and variance, respectively (see Methods). We considered the same normalization and differential methods as in the *muscat* simulation (except MM and *scDD-perm*, which were not considered due to the high computational cost and low statistical power displayed above). As expected, for all methods, differential patterns are easier to detect as the magnitude of the difference increases, with differential location patterns having a higher true positive rate than differential scale patterns. While all methods control the FDR, in all simulations, *distinct* achieves substantially higher TPR than competitors (Figure 6). We also repeated the same simulations including a batch effect, with two batches, with the same scale and location differential parameters for the batch and group differences (i.e., increasing together from 0.2 to 1.5). Again, we excluded *scDD* from these analyses because it cannot handle covariates directly. Results agree with those from the *muscat* batch effect simulation study: FDR and TPRs are mostly unchanged when introducing nuisance covariates, with only a minor decrease in the TPR in stronger batch effects, i.e., when de.facLoc and de.facScale are 1 and 1.5 (Supplementary Figure 8), which again indicates that *distinct* can effectively control for nuisance covariates.

**Figure 6:**
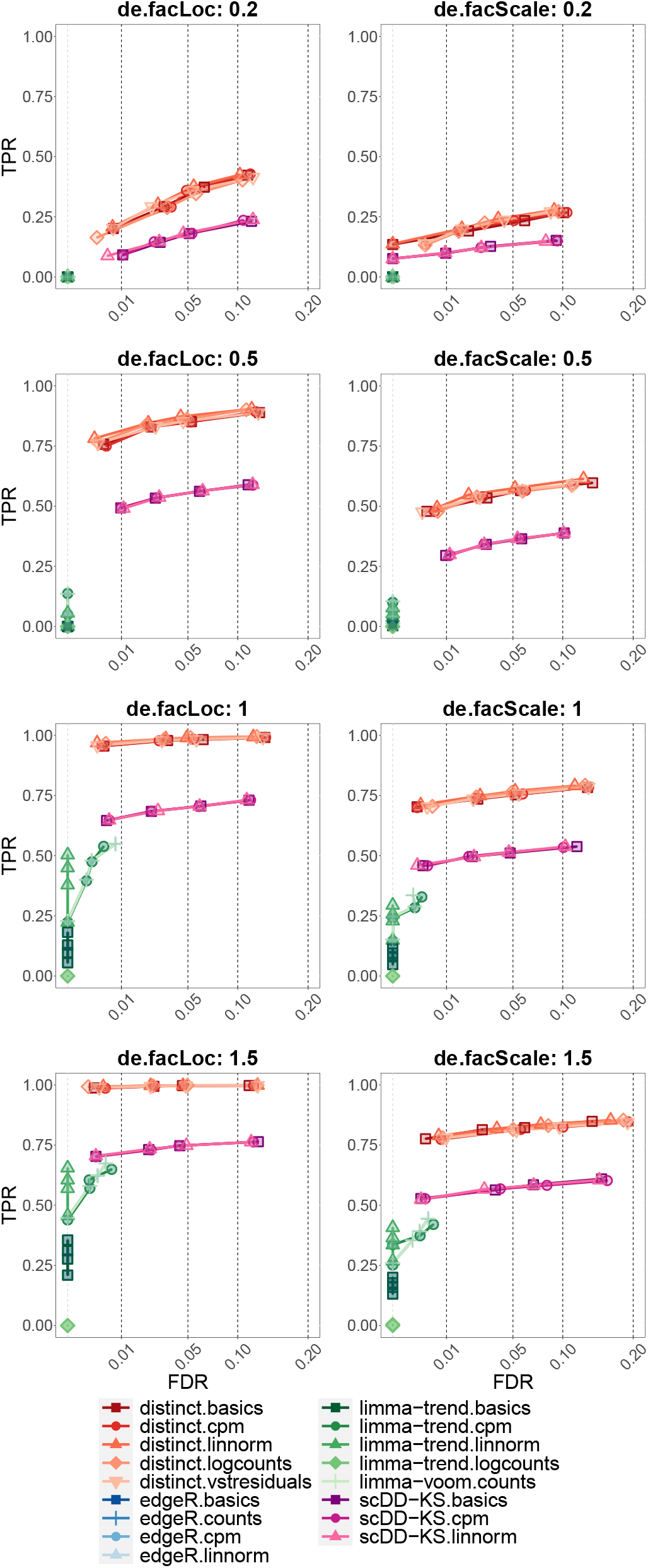
*distinct* displays higher TPR than competitors. TPR *vs*. FDR in *SplatPop* simulated data, with various degrees of differential location (left) and scale (right) parameters, primarily affecting the mean and variance, respectively. Circles indicate observed FDR for 0.01, 0.05, 0.1 and 0.2 significance thresholds. Each simulation consists of 20,345 genes genes, 800 cells (belonging to the same cluster), and two groups of 4 samples each, corresponding to an average of 100 cells per sample.

We further considered the semi-simulated mass cytometry data from Weber *et al*. [31] (labelled *diffcyt* simulation), where spike-in signals were computationally introduced in experimental data [4], hence maintaining the properties of real biological data while also embedding a known ground truth signal. We evaluated *distinct* and two methods from *diffcyt*, based on *limma* [22] and linear mixed models (LMM), which out-performed competitors on these same data [31]. In particular, we considered three datasets from Weber *et al*. [31]: the main DS dataset and two more where differential effects were diluted by 50 and 75%. Each dataset consists of 24 protein markers, 88,435 cells, and two groups (with and without spike-in signal) of 8 samples each. Measurements were first transformed, and then cells were grouped into sub-populations with two separate approaches (see Methods): i) similarly to the *muscat* simulation study, cell labels were defined based on 8 manually annotated cell types [31] (Figure 7a), and ii) as in the original *diffcyt* study from Weber *et al*. [31], cells were grouped into 100 high-resolution clusters (based on 10 cell-type markers, see Methods) via unsupervised clustering (Figure 7b). In the main simulation, *distinct* achieves higher TPR when considering cell-type labels (Figure 7a, ‘main’), while all methods exhibit substantially overlapping performance when using unsupervised clustering (Figure 7b, ‘main’). In both clustering approaches, as the magnitude of the differential effect decreases, the distance between methods increases: *diffcyt* tools show a significant drop in the true positive rate whereas *distinct* maintains a higher TPR while effectively controlling for the false discovery rate (FDR) (Figures 7a-b and Supplementary Figure 9). This indicates that *distinct* has good statistical power to detect even small changes between conditions. We also considered the three replicate null datasets from

**Figure 7:**
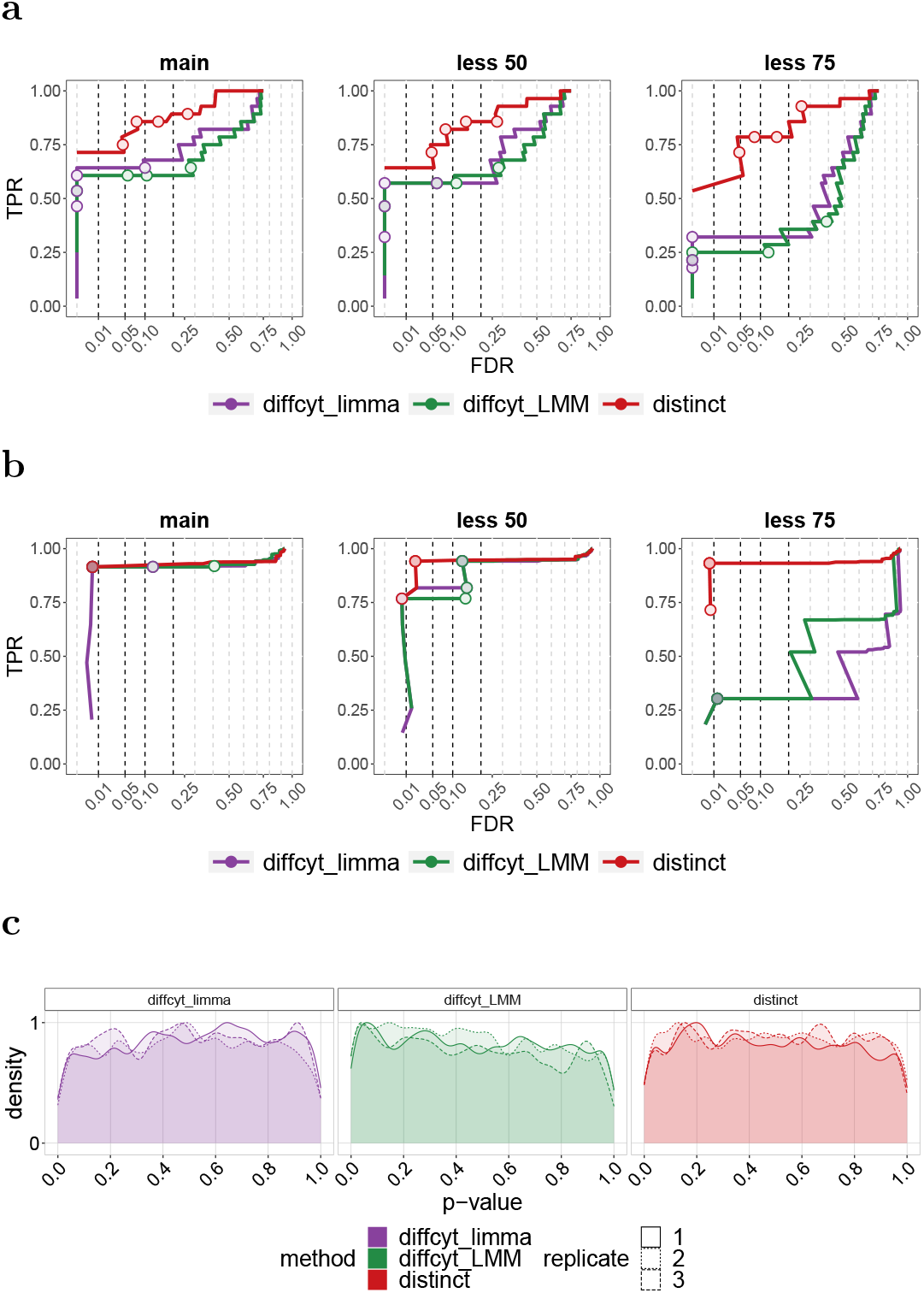
*distinct* shows high power while controlling for false positive and false discovery rates. **(a-b)** TPR *vs*. FDR in *diffcyt* semi-simulated data. ‘main’, ‘less 50’ and ‘less 75’ indicate the main simulation, and those where differential effects are diluted by 50 and 75%, respectively. Each simulation consists of 88,435 cells and two groups of 8 samples each. Circles indicate observed FDR for 0.01, 0.05, 0.1 and 0.2 significance thresholds. **(a)** As in the *muscat* simulation study, cells were clustered into 8 populations based on manually annotated cell types [31]. **(b)** As in Weber *et al*. [31], cells were grouped in 100 high-resolution clusters via unsupervised clustering. **(c)** Density of raw p-values in *diffcyt* null semi-simulated data; each replicate represents a different null simulation. Each replicate consists of 88,438 cells and two groups of 8 samples each. As in Weber *et al*. [31], cells were clustered in an unsupervised manner.

Weber *et al*. [31] (i.e., with no differential effect), containing 24 protein markers and 88,438 cells across 8 cell types, and found that all methods display approximately uniform p-values (Figure 7c).

### Experimental data analyses

In order to investigate false positive rates (FPRs) in real data, we considered two experimental scRNA-seq datasets where no differential signals were expected, by comparing samples from the same experimental condition. Given the high computational cost and low power of MM, and the high FDR of *scDD* models, for the real data analyses, we only included *distinct* and PB methods. We considered gene-cluster combinations with at least 20 non-zero cells across all samples. The first dataset (labelled *T-cells*) consists of a Smart-seq2 scRNA-seq dataset of 19,875 genes and 11,138 T cells isolated from peripheral blood from 12 colorectal cancer patients [35]. We automatically separated cells in 11 clusters (via *igraph* [1, 7]), and generated replicate datasets, by randomly separating, three times, the 12 patients to two groups of size 6. The second dataset (labelled *Kang*) contains 10x droplet-based scRNA-seq peripheral blood mononuclear cell data from 8 Lupus patients, before (controls) and after (stimulated) 6h-treatment with interferon-*β* (INF-*β*), a cytokine known to alter the transcriptional profile of immune cells [12]. The full dataset contains 35,635 genes and 29,065 cells, which are separated (via manual annotation [12]) into 8 cell types. One of the 8 patients was removed as it appears to be a potential outlier (Supplementary Figures 10-12). Here we only included singlet cells and cells assigned to a cell population, and considered control samples only, resulting in 11,854 cells and 10,891 genes. Again, we artificially created three replicate datasets by randomly assigning the 7 retained control samples in two groups of size 3 and 4. In both null analyses, we found that *limma-trend*, particularly when using CPMs, leads to an increase of FPRs, *distinct* ‘s p-values are only marginally inflated towards 0, while *edgeR* and *limmavoom* are the most conservative methods and provide the best control of FPRs (Figure 8 and Supplementary Tables 2-3). Regarding normalization, *linnorm* and *BASiCS* lead to the most conservative p-values and smallest false positive rates.

**Figure 8:**
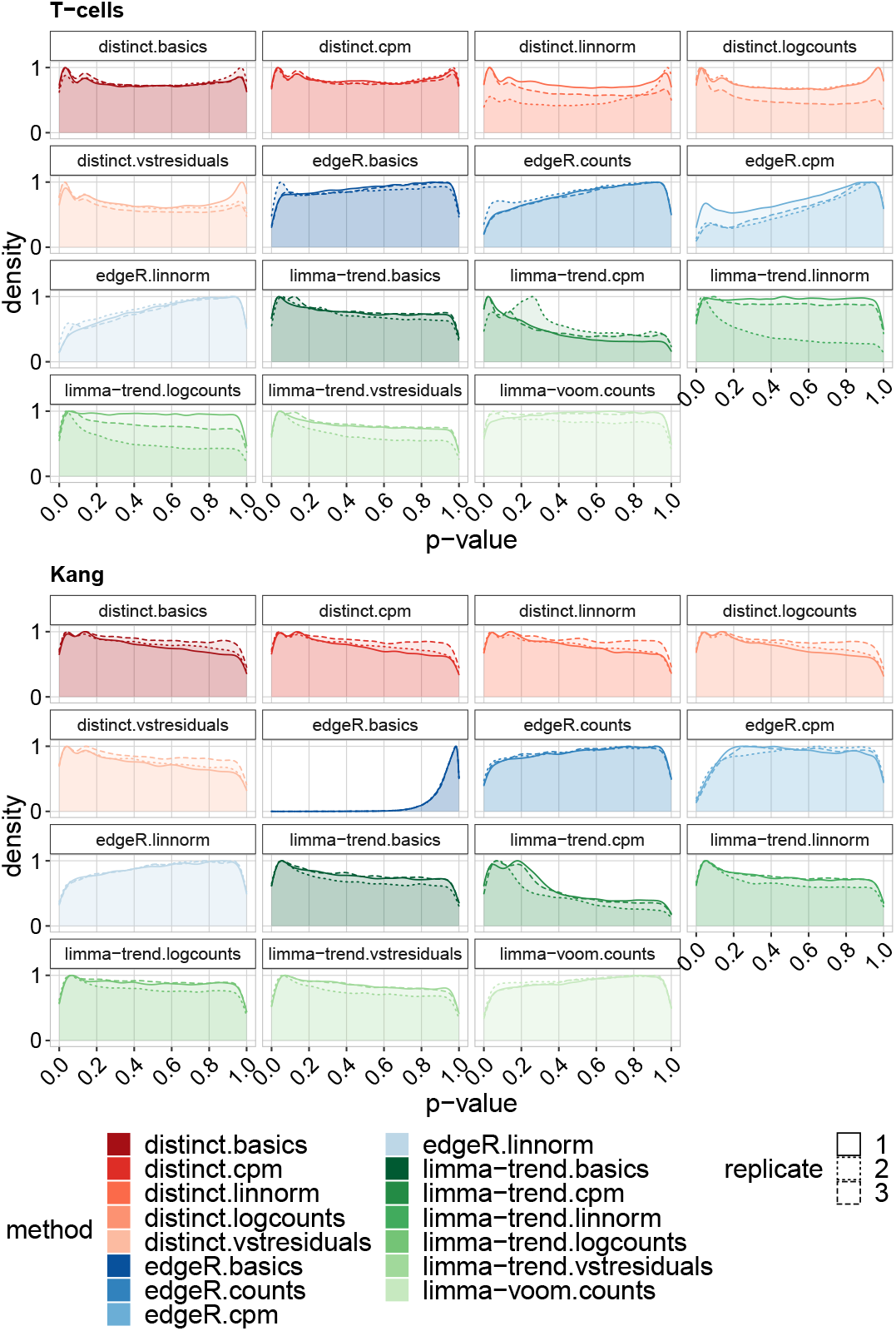
On experimental scRNA-seq data, *distinct* has almost-uniform null p-values. Density of raw p-values in the null *T-cells* (top) and *Kang* (bottom) experimental data. Each replicate represents a random partition of samples in two groups. The *T-cells* data consists of 12 samples and 11,138 cells across 11 clusters. For the *Kang* dataset, we retained 7 samples and 11,854 cells across 8 clusters.

We then considered again the *Kang* dataset, and performed a DS analysis between controls and stimulated samples. Again, we removed one potential outlier patient, and only considered singlet cells and cells assigned to a cell population; we further filtered genecluster combinations with less than 20 non-zero cells across all samples, resulting in 12,045 genes and 23,571 cells across 8 cell types and 14 samples. We found that *distinct* identifies more differential patterns than PB methods, with *edgeR* and *limma-voom* being the most conservative methods, and that its results are very coherent across different input data (Supplementary Figure 13). When visually investigating the gene-cluster combinations detected by *distinct* (adjusted p-value < 0.1), on all five input data (CPMs, logcounts, linnorm, BASiCS and vstresiduals), and not detected by any of the ten PB approaches (adjusted p-value > 0.1), we found several interesting non-canonical differential patterns (Figure 9 and Supplementary Figures 14-25). In particular, gene MARCKSL1 displays a DB pattern, with stimulated samples having higher density on the tails and lower in the centre of the distribution, gene RPL13 mirrors classical DE, while the other genes seem to emulate DP profiles. Interestingly, ten out of eleven of these genes are known tumor prognostic markers: H2AZ2 for cervical and renal cancer, SRSF9 for liver cancer and melanoma, RPL24 for renal and thyroid cancer, HNRNPA0 for renal and pancreatic cancer, MARCKSL1 for liver and renal cancer, GTF3C6 for liver cancer, RPL13 for endometrial and renal cancer, PGK1 for breast, head and neck, cervical, liver, and pancreatic cancer, KDELR2 for renal, head and neck and glioma cancer, and RPL11 for renal and breast cancer [27]. This is an interesting association, considering that INF-*β* stimulation is known to inhibit and interfere with tumor progression [8,21]. Additionally, Supplementary Figures 9-17 show how *distinct* can identify differences between groups of distributions even when only a portion of the ECDF varies between conditions. Finally, we computed the fraction of detected genes that are unique by each method. Given that a ground truth is absent, we speculate that gene-cluster combinations detected by multiple methods are more likely to be truly differential, while those detected by a single method are more likely to be false positive detections. Since methods return widely different number of significant genes, for each method, we considered the top (i.e., smallest p-value) 1,000 genes per cell-type. We then computed the percentage of results that are unique to each method (Table 1), i.e., not in common with the top 1,000 results returned by any other method. Overall, *distinct* displays a lower fraction of unique results (1.4% on average across all input data) compared to *edgeR* (4%) and *limma* (6.7%). It is also interesting to note that *scater* ‘s logcounts normalization lead to the 2 smallest fractions of unique values (i.e., *distinct*.*logcounts* and *limma-trend*.*logcounts*).

**Figure 9:**
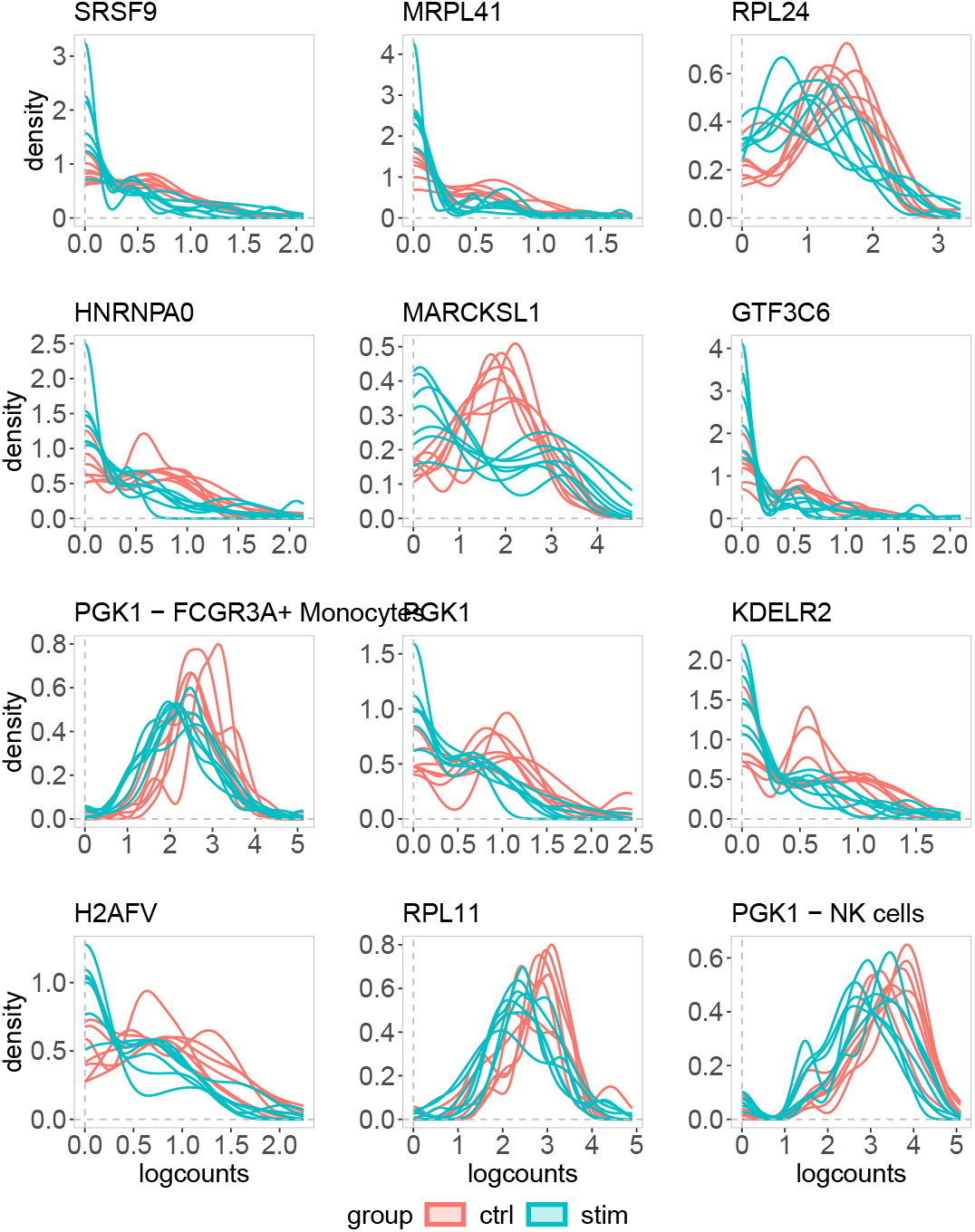
*distinct* discovers non-canonical differential patterns. Density of logcounts for nine examples of differential patterns identified by *distinct* on all input data (adjusted p-values < 0.05), and not by any PB tool (adjusted p-values > 0.05), on the *Kang* dataset when comparing controls and stimulated samples. Gene RPL13 was identified in FCGR3A+ Monocytes (third row) and in NK cells (fourth row), while all other genes were detected in Dendritic cells. Each line represents a sample.

## Discussion

High-throughput single-cell data can display complex differential patterns; nonetheless, most methods for differential expression fail to identify changes where the mean is not affected. To overcome the limitations of present differential tools, we have developed *distinct*, a novel method to identify differential patterns between groups of distributions, which is particularly well suited to perform differential analyses on high-throughput single-cell data. *distinct* is based on a flexible hierarchical multi-sample full-distribution non-parametric approach. In order to compare it to state-of-the-art differential methods, we ran extensive benchmarks on both simulated and experimental datasets from scRNA-seq and mass cytometry data, where our approach exhibits favourable performance, provides good control of the FNR and FDR, and is able to identify more patterns of differential expression compared to canonical tools, even when the overall mean is unchanged. In particular, our approach displays a higher statistical power (i.e., TPR) not only than PB methods, but also compared to other non-parametric frameworks from *scDD*, based on the Kolmogorov-Smirnov test statistic (*scDD-KS*) and on permutation tests (*scDD-perm*). *distinct* also allows for biological replicates, does not rely on asymptotic theory, which could be inaccurate in small sample sizes (typical of biological data), and avoids parametric assumptions, that may be challenging to meet in single-cell data. Additionally, *distinct* can also effectively adjust for sample-level cell-cluster specific covariates (i.e., whose effect varies across cell clusters), such as batch effects (Supplementary Figure 3). Importantly, *distinct* is a very general test that, due to its non-parametric nature, can be applied to various types of data, even beyond the single-cell applications shown here. Furthermore, thanks to its flexible form, we have shown in our simulations that *distinct* has the most consistent performance across normalization approaches (Supplementary Figure 2).

However, these advantages come at the expense of a higher computational burden, particularly when compared to PB methods or KS approaches (Figure 5). Nonetheless, by employing clever computational techniques (i.e., parallel computing and C++ coding within R), the method runs within minutes on a laptop, even for large datasets. Overall, we believe that *distinct* represents a valid alternative for differential detections from single-cell data, particularly when interest lies beyond canonical differences in means, as it allows to enhance statistical power at the cost of a reasonable increase in the computational time.

Finally, although we have focused here on comparing two groups of samples, several future extensions are possible to allow our framework to be applied to different scenarios. For instance, by suitably modifying the test statistics in (1), one may ideally extend our approach to perform a joint differential test between three of more groups of samples. Although, it is worth noting that, in the presence of three or more experimental conditions, at present, it is still possible to run pairwise comparisons between pairs of conditions. While a joint test across all groups may certainly be of interest in some cases, from our experience, comparisons between pairs of groups are usually more used among scientists. In addition, as we were suggested by a user, *distinct* could be employed to compare cell clusters instead of experimental conditions, hence discovering differential genes between cell clusters (e.g., cell types), even from individual samples.

## Availability

*distinct* is freely available as a Bioconductor R package at: https://bioconductor.org/packages/distinct. The scripts used to run all analyses are available on GitHub (https://github.com/SimoneTiberi/distinct_manuscript, version v3) and Zenodo (DOI: 10.5281/zenodo.6397114). The *diffcyt* simulated data is available via FlowRepository (accession ID FR-FCM-ZYL8 [31]) and *HDCytoData* R Bioconductor package [32]; the *Kang* dataset can be accessed via *musc-Data* R Bioconductor package [5]; the *T-cells* dataset is deposited on the European Genome-phenome (accession id EGAD00001003910 [35]).

## Acknowledgements

We acknowledge Almut Luetge, Brian D M Tom, Christina Azodi, Davis McCarthy, Reinhard Furrer, and the entire Robinson lab for precious comments and suggestions. This work was supported by Forschungskredit to ST (grant number FK-19-113) as well as by the Swiss National Science Foundation to MDR (grants 310030_175841, CRSII5_177208). MDR acknowledges support from the University Research Priority Program Evolution in Action at the University of Zurich.

## Author contributions

ST conceived the method, implemented it, performed all analyses and wrote the manuscript. ST and MDR designed the study. HLC and LMW contributed to *muscat* and *diffcyt* simulation studies, respectively. PS contributed to the computational development of *distinct* and to the revision process. All authors read, contributed to, and approved the final article.

## Methods

### Permutation test

In order to test for differences between groups, we employ a hierarchical permutation approach: to estimate the null distribution of *s*^*obs*^, we permute the individual observations (e.g., single-cell measurements) instead of the samples. Note that this violates the exchangeability assumption of permutation tests and, hence, p-values are not guaranteed to be uniformly distributed under the null hypothesis; nonetheless, in our simulated and experimental analyses, we empirically show that *distinct* provides good control of both false positive and false discovery rates. We randomly permute individual observations *P* times across all samples and groups, by retaining the original sample sizes. We denote by *s*_*p*_ the test statistic computed at the *p*-th permutation, *p* = 1, …, *P*. A p-value, 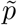, is obtained as [20]:

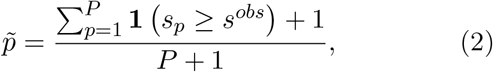

where **1**(*cond*) is 1 if *cond* is true, and 0 otherwise. In order to accurately infer small p-values, when 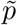 is below some pre-defined thresholds, the number of permutations are automatically increased and 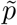 is re-computed. By default, *distinct* initially computes 100 permutations; when 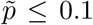 these are increased to 500; when the new 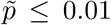we use 2, 000 permutations, which are further increased to 10, 000 if 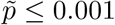. Note that the number of permutations (i.e., 100, 500, 2,000 and 10,000) can be specified by the user.

### Covariates

Assume we observe *Z* nuisance covariates, and that *N* samples are available across all groups, where for the *i*-th sample we observe *C*_*i*_ values (e.g., single-cell measurements). We fit the following linear mixed effects model:

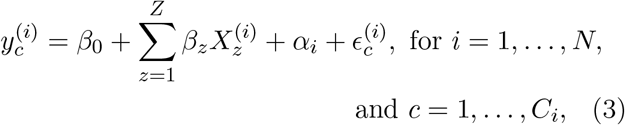

where 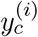 represents the *c*-th observation for the *i*-th sample, *β*_0_ is the intercept of the model, 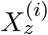 indicates the *z*-th covariate in the *i*-th sample, *β*_*z*_ denotes the fixed effect coefficient for the *z*-th covariate, *α*_*i*_ represents the random effect term for the *i*-th sample, and 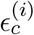 is the (zero-mean) residual for the *c*-th observation in the *i*-th sample. We assume that random terms are normally distributed as 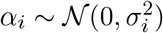, where 𝒩 (*a, b*) denotes the normal distribution with mean *a* and variance *b*. Note that, due to the random effect terms, observations from the same sample are positively correlated while, observations between different samples are independent. We infer model parameters via maximum likelihood, with the estimated values for the fixed effect terms denoted by 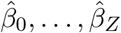. We then remove the estimated effect of nuisance covariates as 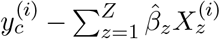; differential testing is performed, as described above, on these normalized values. In DS analyses, model (3) is fit, separately, for every genecluster combination, hence accommodating for cell-type specific effects of covariates.

### Normalization

In scRNA-seq datasets, CPMs and logcounts were computed via *scater* Bioconductor R package [18], vstresiduals were calculated via *sctransform* R package [11] (except for the *T-cells* data, where, due to a failure of *sctransform*’s variance stabilizing normalization, we used *DESeq2* ‘s vst transformation [15]), while *linnorm* and *BASiCS* normalized data were calculated with the respective Bioconductor R packages [9, 28, 29, 33].

In mass cytometry datasets, measurements were transformed via *diffcyt* ‘s *transformData* function, which applies an *arcsinh* transformation.

### *muscat* simulation and *Kang* data

In all *muscat* simulations, we used the control samples of the *Kang* dataset as a anchor data; as in the real data analyses, we excluded one sample as it emerged as a potential outlier (Supplementary Figures 5-7), and only considered singlet cells and cells assigned to a cell population. In *muscat* ‘s simulation studies, we considered gene-cluster combinations with simulated expression mean greater than 0.2; for DB patterns, we increased this threshold to 1 because with low expression values differences are not visible by eye. In the simulation when varying the library size (Supplementary Figure 4), we filtered gene-clusters combinations with at least 50 non-zero cells. For every simulations, five replicates were simulated, and results were averaged across replicates. In the main simulation (Figure 2) and the batch effect simulation (Supplementary Figure 3), we simulated from a paired design 2 groups of 3 samples each, with 4,000 genes, and 3,600 cells distributed in 3 clusters (corresponding to an average of 200 cells per sample in each cluster). For the simulation study when varying the number of cells (Figure 4 and Supplementary Figure 3), the total numbers of available cells were 900, 1,800, 3,600 and 7,200, corresponding to an average of 50, 100, 200 and 400 cells per sample in every cluster. For the differential simulations, we used log2-FC values of 1 for DE, 1.5 for DP and DM, and 3 for DB and DV. For the batch effect simulation study we used a modified version of *muscat*, developed by Almut Luetge at the Robinson lab (available at: https://github.com/SimoneTiberi/distinct_manuscript), which allows simulating cluster-specific batch effects [6, 16]. All *muscat* simulation studies, as well as the *Kang* non-null data analysis, were performed by editing the original snakemake workflow from Crowell *et al*. [6]. PB methods were applied on aggregated data by summing cell-level measurements; for differential testing, we used *muscat* ‘s *pbDS* function [6]. Mixed model methods were implemented, via *muscat* ‘s *mmDS* function, using the same approaches as in Crowell *et al*. [6]: in *MM-dream2* and *MM-vstresiduals* linear mixed models were applied to log-normalized data with observational weights and variance-stabilized data, respectively, while in *MM-nbinom* generalized linear mixed models were fitted directly to raw counts. In the *muscat* simulations and in the *Kang* non-null data analysis, we accounted for the paired design by modelling the patient id as a covariate in all methods that allow for covariates (i.e., *distinct*, PB and MM).

### *splatPop* simulation

In *SplatPOP* simulated data, we used a human genome, version 19, downloaded from https://www.gencodegenes.org/human/release_19.html. We ran a total of 16 simulations: 8 with and 8 without batch effects as nuisance covariate. In each case, we ran 4 differential location (“de.facLoc” parameter) and 4 differential scale (“de.facScale” parameter) simulations, with differential parameters equals to 0.2, 0.5, 1 and 1.5. In every simulation, 10% of genes were differential between groups, and a total of 20,345 genes and 800 cells were simulated (100 per sample). In the simulation with batch effects, the 8 samples were randomly assigned to 2 batches, and the differential location and scale parameters between batches (“batch.facLoc” and “batch.facScale”, respectively) matched those between groups of samples (“de.facLoc” and “de.facScale”). For more details on how *SplatPOP* ‘s data is simulated, please refer to the original manuscript [2] and vignettes.

### *diffcyt* simulation

The *diffcyt* semi-simulated data originates from a real mass cytometry dataset of healthy peripheral blood mononuclear cells from two paired groups of 8 samples each [4]; one group contains unstimulated cells, while the other was stimulated with B cell receptor/Fc receptor cross-linker. The original dataset contains a total of 172,791 cells and 24 protein markers: 10 of these are cell-type markers used for cell clustering, while 14 are cell state markers used for differential state analyses; the distinction between cell state and cell-type markers is based on prior biological knowledge [31]. In Weber *et al*. [31], semi-simulated data were generated by separating the cells of each unstimulated sample in two artificial samples; a differential signal was then computationally introduced by replacing, in one group, unstimulated B cells with B cells from stimulated samples. Measurements were transformed and cells clustered via *diffcyt* ‘s *transformData* (which applies an *arcsinh* transformation) and *generateClusters* functions, respectively. For the DS simulation in Figure 7b, as in Weber *et al*. [31], we evaluated methods’ performance in terms of detecting DS for phosphorylated ribosomal protein S6 (pS6) in B cells, which is the strongest differential signal across the cell types in this dataset [19, 31]. For the DS simulation in Figure 7a, we considered previously manually annotated cell types [31] and included all 14 cell state markers. *diffcyt* ‘s *limma* and LMM methods were applied via *diffcyt* ‘s *testDS_limma* and *testDS_LMM* functions, respectively [31]. We accounted for the paired design by modelling the patient id as a covariate.

### P-values adjustment

All p-values were adjusted via Benjamini-Hochberg correction [3]. In *diffcyt* simulations we used globally adjusted p-values for all methods, i.e., p-values from all clusters are jointly adjusted once. However, since PB methods were found to be over-conservative when globally adjusting p-values [6], in *muscat* simulations and *Kang* discovery analyses, we used locally adjusted p-values for all methods.

### Software versions

All analyses were performed via R software version 4.0.0, with Bioconductor packages from release 3.11.

