## Supplementary material for "*distinct*: a novel approach to differential distribution analyses"

#### 1 Supplementary Tables

| | DE | DP | DM | DB | DV | null | average |
|---|---|---|---|---|---|---|---|
| scDD-perm.vstresiduals | 2191.8 | 2083.2 | 2053.2 | 1631.9 | 2244.0 | 2309.2 | 2085.6 |
| scDD-perm.linnorm | 593.4 | 574.9 | 591.9 | 662.8 | 587.0 | 552.2 | 593.7 |
| scDD-perm.cpm | 531.5 | 541.4 | 548.9 | 592.4 | 541.8 | 512.9 | 544.8 |
| scDD-perm.basics | 534.7 | 541.9 | 553.2 | 580.1 | 545.0 | 513.0 | 544.7 |
| MM-nbinom | 318.0 | 314.7 | 322.0 | 298.2 | 207.2 | 323.7 | 297.3 |
| MM-vstresiduals | 85.5 | 90.8 | 87.8 | 95.0 | 66.8 | 81.5 | 84.6 |
| MM-dream2 | 30.3 | 30.7 | 28.6 | 31.4 | 25.2 | 30.4 | 29.4 |
| distinct.vstresiduals | 6.7 | 6.0 | 5.9 | 3.7 | 2.7 | 2.0 | 4.5 |
| distinct.logcounts | 5.3 | 4.4 | 4.5 | 3.2 | 2.2 | 1.5 | 3.5 |
| distinct.cpm | 5.2 | 4.5 | 4.5 | 3.0 | 1.9 | 1.5 | 3.4 |
| distinct.linnorm | 4.8 | 4.3 | 4.5 | 3.2 | 2.1 | 1.5 | 3.4 |
| distinct.basics | 4.6 | 4.3 | 4.5 | 2.6 | 1.9 | 1.5 | 3.2 |
| scDD-KS.vstresiduals | 0.7 | 0.7 | 0.7 | 0.9 | 0.6 | 0.6 | 0.7 |
| scDD-KS.cpm | 0.5 | 0.5 | 0.5 | 0.5 | 0.5 | 0.5 | 0.5 |
| scDD-KS.linnorm | 0.5 | 0.5 | 0.5 | 0.6 | 0.4 | 0.4 | 0.5 |
| scDD-KS.basics | 0.4 | 0.4 | 0.5 | 0.5 | 0.4 | 0.4 | 0.5 |
| edgeR.cpm | 0.2 | 0.2 | 0.2 | 0.2 | 0.2 | 0.2 | 0.2 |
| edgeR.counts | 0.2 | 0.2 | 0.2 | 0.2 | 0.2 | 0.1 | 0.2 |
| edgeR.linnorm | 0.2 | 0.2 | 0.2 | 0.2 | 0.2 | 0.2 | 0.2 |
| edgeR.basics | 0.1 | 0.1 | 0.1 | 0.1 | 0.1 | 0.1 | 0.1 |
| limma-trend.cpm | 0.1 | 0.1 | 0.1 | 0.1 | 0.1 | 0.1 | 0.1 |
| limma-voom.counts | 0.1 | 0.1 | 0.1 | 0.1 | 0.1 | 0.1 | 0.1 |
| limma-trend.basics | 0.1 | 0.1 | 0.1 | 0.1 | 0.1 | 0.1 | 0.1 |
| limma-trend.linnorm | 0.1 | 0.1 | 0.1 | 0.1 | 0.1 | 0.1 | 0.1 |
| limma-trend.vstresiduals | 0.1 | 0.1 | 0.1 | 0.1 | 0.1 | 0.1 | 0.1 |
| limma-trend.logcounts | 0.1 | 0.1 | 0.1 | 0.1 | 0.1 | 0.1 | 0.1 |

**Supplementary Table 1:** Computing time, expressed in minutes, in *muscat* main simulations. For each column, times are averaged across the five replicate simulations; in each replicate simulation 3,600 cells are available (200, on average, per cluster-sample combination). 'MM' refers to mixed models, *scDD-KS* indicates *scDD* based on the Kolmogorov-Smirnov test, while *scDD-perm* denotes *scDD* based on the permutation test (100 permutations). *distinct*, MM models and *scDD* were run on 3 cores, while pseudo-bulk methods based on *edgeR* and *limma* used a single core since they do not allow for parellel computing. Note that computing times for *distinct* are significantly larger in non-null datasets (which contain, on average, 10% of differential genes in each cluster), because *distinct* increases the number of permutations used (and hence the computational cost) for significant results. Note that *scDD-perm* requires much longer on vstresiduals than on the other normalized data, because *scDD* performs differential testing on non-zero values: vstresiduals, (unlike linnorm, cpm and basics normalized data) are not zero-inflated and, therefore, many more cells have to be used for differential testing.

| threshold | raw p-value | | | globally adjusted p-value | | | locally adjusted p-value | | |
|---|---|---|---|---|---|---|---|---|---|
| | 0.1 | 0.05 | 0.01 | 0.1 | 0.05 | 0.01 | 0.1 | 0.05 | 0.01 |
| distinct.basics | 0.12 | 0.08 | 0.02 | 0.01 | 0.01 | 0.00 | 0.01 | 0.01 | 0.00 |
| distinct.cpm | 0.12 | 0.08 | 0.02 | 0.01 | 0.00 | 0.00 | 0.01 | 0.01 | 0.00 |
| distinct.linnorm | 0.13 | 0.09 | 0.03 | 0.01 | 0.01 | 0.00 | 0.01 | 0.01 | 0.00 |
| distinct.logcounts | 0.15 | 0.10 | 0.04 | 0.02 | 0.01 | 0.00 | 0.02 | 0.01 | 0.00 |
| distinct.vstresiduals | 0.15 | 0.10 | 0.03 | 0.01 | 0.01 | 0.00 | 0.01 | 0.01 | 0.00 |
| edgeR.basics | 0.09 | 0.05 | 0.01 | 0.00 | 0.00 | 0.00 | 0.00 | 0.00 | 0.00 |
| edgeR.counts | 0.07 | 0.03 | 0.01 | 0.00 | 0.00 | 0.00 | 0.00 | 0.00 | 0.00 |
| edgeR.cpm | 0.07 | 0.03 | 0.00 | 0.00 | 0.00 | 0.00 | 0.00 | 0.00 | 0.00 |
| edgeR.linnorm | 0.06 | 0.03 | 0.00 | 0.00 | 0.00 | 0.00 | 0.00 | 0.00 | 0.00 |
| limma-trend.basics | 0.14 | 0.08 | 0.02 | 0.00 | 0.00 | 0.00 | 0.01 | 0.00 | 0.00 |
| limma-trend.cpm | 0.19 | 0.12 | 0.05 | 0.00 | 0.00 | 0.00 | 0.04 | 0.04 | 0.00 |
| limma-trend.linnorm | 0.15 | 0.09 | 0.03 | 0.00 | 0.00 | 0.00 | 0.02 | 0.01 | 0.00 |
| limma-trend.logcounts | 0.14 | 0.08 | 0.02 | 0.00 | 0.00 | 0.00 | 0.01 | 0.01 | 0.00 |
| limma-trend.vstresiduals | 0.14 | 0.08 | 0.02 | 0.00 | 0.00 | 0.00 | 0.01 | 0.01 | 0.00 |
| limma-voom.counts | 0.11 | 0.06 | 0.02 | 0.00 | 0.00 | 0.00 | 0.01 | 0.01 | 0.00 |

**Supplementary Table 2:** Fraction of false positive tests returned from each method, at the 0.01, 0.05 and 0.1 significance thresholds, in the null *T-cells* experimental data. Values are averages across the three replicates.

| threshold | raw p-value | | | globally adjusted p-value | | | locally adjusted p-value | | |
|---|---|---|---|---|---|---|---|---|---|
| | 0.1 | 0.05 | 0.01 | 0.1 | 0.05 | 0.01 | 0.1 | 0.05 | 0.01 |
| distinct.basics | 0.13 | 0.08 | 0.02 | 0.01 | 0.00 | 0.00 | 0.01 | 0.01 | 0.00 |
| distinct.cpm | 0.13 | 0.08 | 0.02 | 0.01 | 0.00 | 0.00 | 0.01 | 0.01 | 0.00 |
| distinct.linnorm | 0.13 | 0.08 | 0.02 | 0.01 | 0.00 | 0.00 | 0.01 | 0.01 | 0.00 |
| distinct.logcounts | 0.13 | 0.08 | 0.03 | 0.01 | 0.01 | 0.00 | 0.01 | 0.01 | 0.00 |
| distinct.vstresiduals | 0.14 | 0.09 | 0.03 | 0.01 | 0.00 | 0.00 | 0.01 | 0.01 | 0.00 |
| edgeR.basics | 0.00 | 0.00 | 0.00 | 0.00 | 0.00 | 0.00 | 0.00 | 0.00 | 0.00 |
| edgeR.counts | 0.09 | 0.05 | 0.01 | 0.00 | 0.00 | 0.00 | 0.00 | 0.00 | 0.00 |
| edgeR.cpm | 0.05 | 0.02 | 0.00 | 0.00 | 0.00 | 0.00 | 0.00 | 0.00 | 0.00 |
| edgeR.linnorm | 0.08 | 0.04 | 0.01 | 0.00 | 0.00 | 0.00 | 0.00 | 0.00 | 0.00 |
| limma-trend.basics | 0.14 | 0.08 | 0.02 | 0.00 | 0.00 | 0.00 | 0.00 | 0.00 | 0.00 |
| limma-trend.cpm | 0.20 | 0.11 | 0.03 | 0.00 | 0.00 | 0.00 | 0.02 | 0.01 | 0.00 |
| limma-trend.linnorm | 0.15 | 0.08 | 0.02 | 0.00 | 0.00 | 0.00 | 0.00 | 0.00 | 0.00 |
| limma-trend.logcounts | 0.12 | 0.07 | 0.02 | 0.00 | 0.00 | 0.00 | 0.00 | 0.00 | 0.00 |
| limma-trend.vstresiduals | 0.13 | 0.07 | 0.01 | 0.00 | 0.00 | 0.00 | 0.00 | 0.00 | 0.00 |
| limma-voom.counts | 0.08 | 0.04 | 0.01 | 0.00 | 0.00 | 0.00 | 0.00 | 0.00 | 0.00 |

**Supplementary Table 3:** Fraction of false positive tests returned from each method, at the 0.01, 0.05 and 0.1 significance thresholds, in the null *Kang* experimental data. Values are averages across the three replicates.

### 2 Supplementary Figures

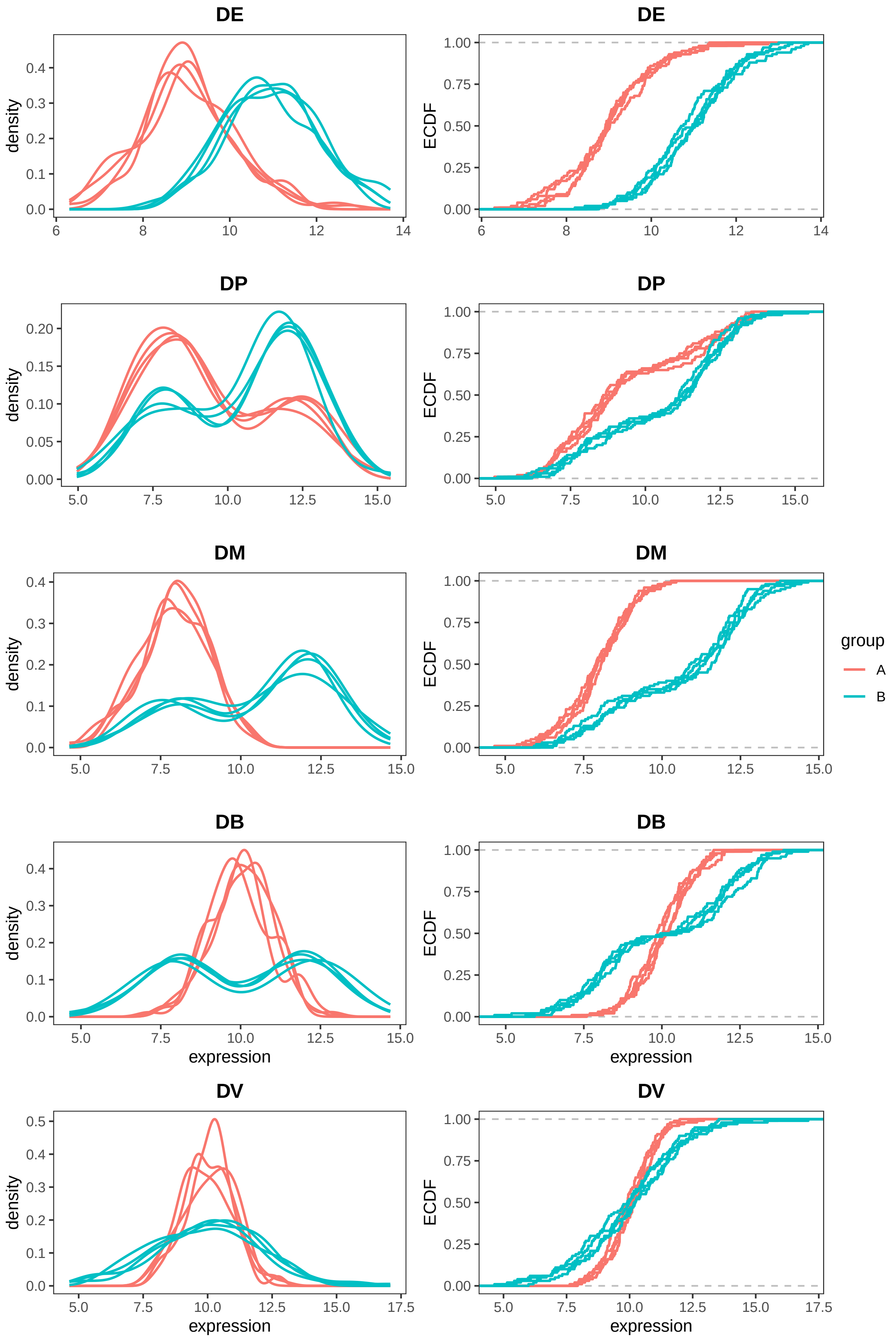


**Supplementary Figure 1:** Density and corresponding empirical cumulative distribution function (ECDF) for differential patterns DE, DP, DM, DB and DV, as illustrated in Figure 1. Two groups of four samples each are compared; 100 cells were simulated from each sample.

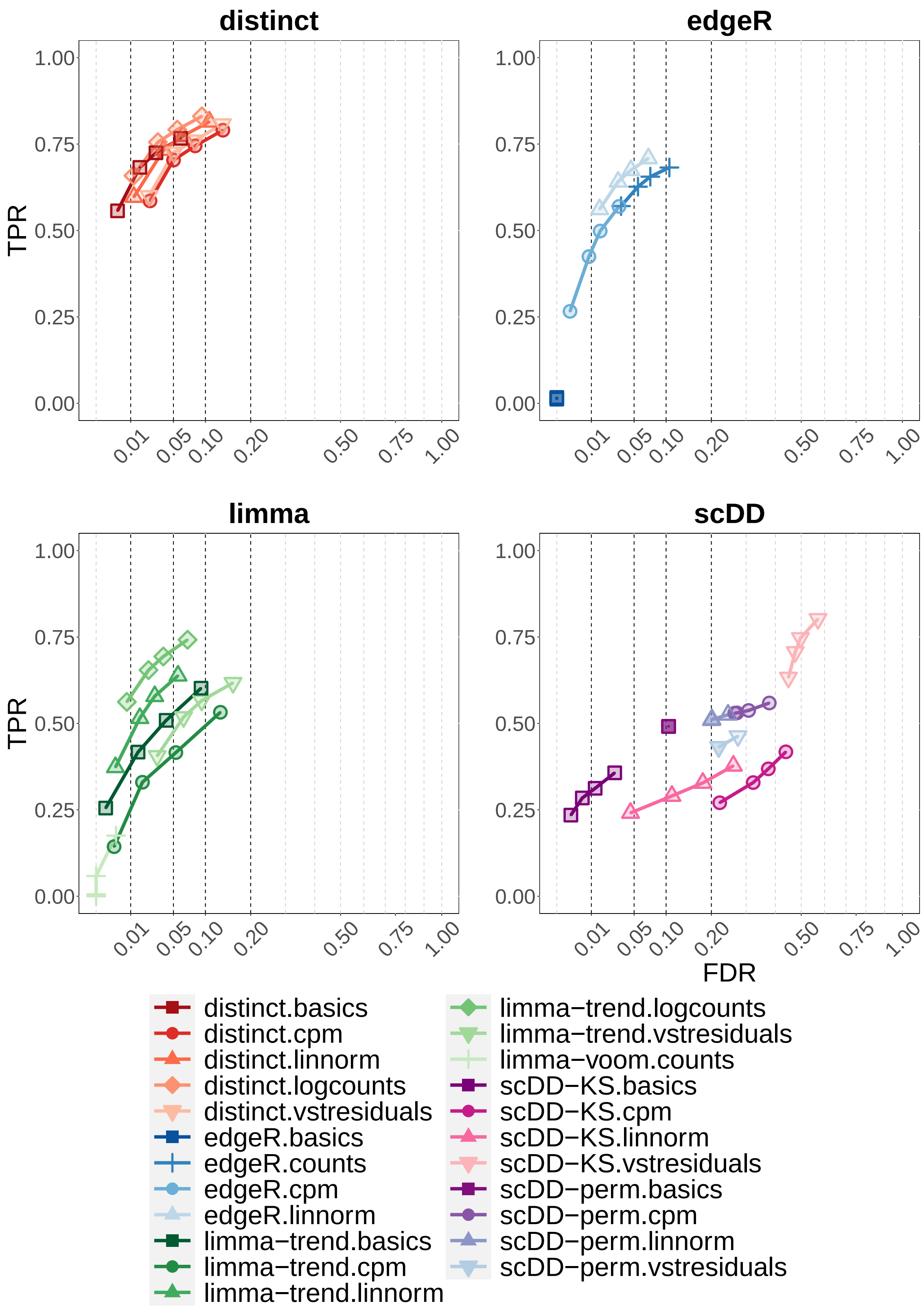


**Supplementary Figure 2:** TPR vs. FDR in *muscat* simulated data with batch effects; DE, DP, DM, DB and DV refer to the differential profiles illustrated in Figure 1. Circles indicate observed FDR for 0.01, 0.05, 0.1 and 0.2 significance thresholds. Results are averages across the five simulation replicates. Each individual replicate consists of 4,000 genes, 3,600 cells, separated into 3 clusters, and two groups of 3 samples each, corresponding to an average of 200 cells per sample in each cluster. Note that *scDD* methods were excluded from this analysis because *scDD* does not directly handle covariates.

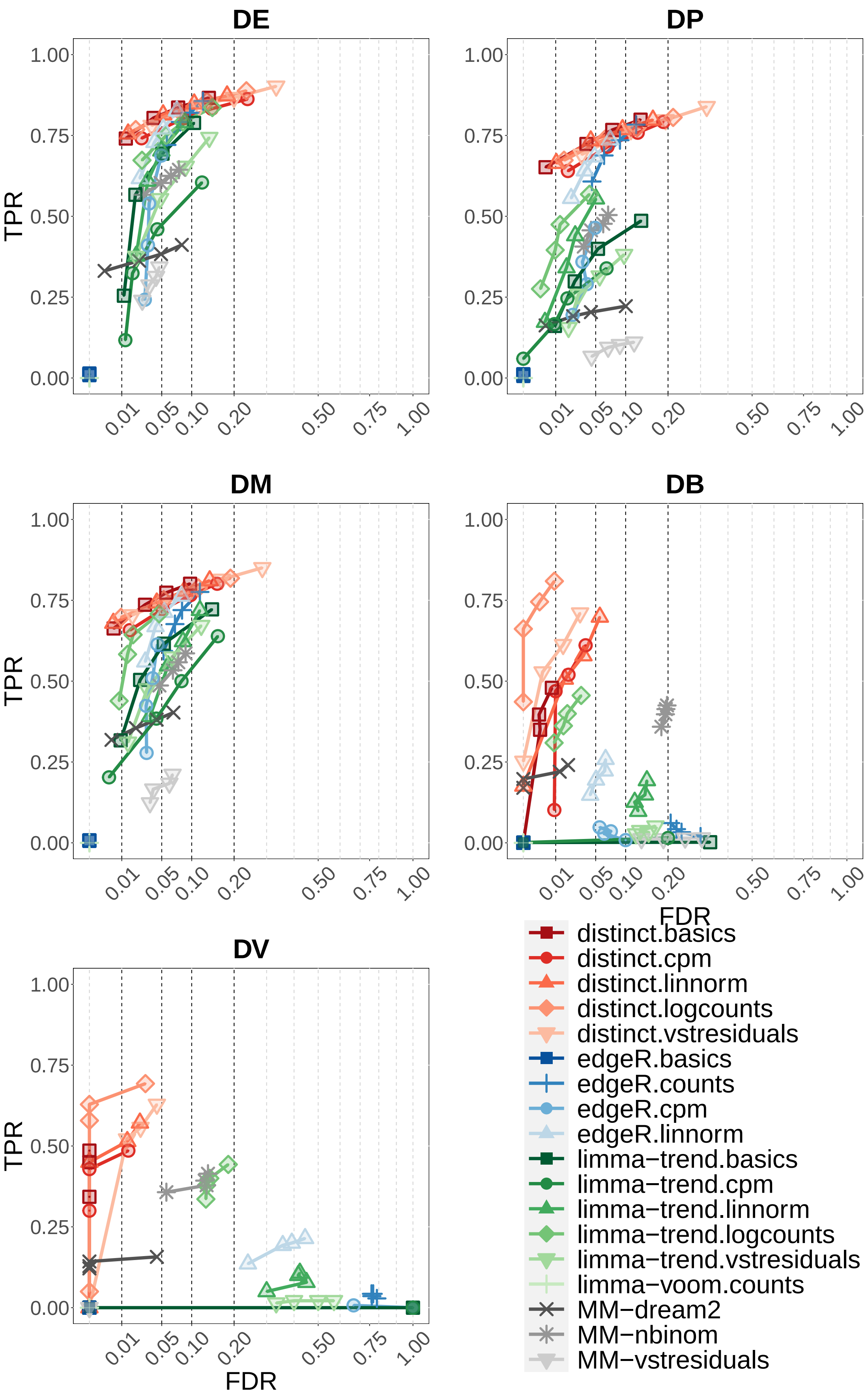


**Supplementary Figure 3:** TPR vs. FDR in *muscat* simulated data with batch effects; DE, DP, DM, DB and DV refer to the differential profiles illustrated in Figure 1. Results are averages across the five simulation replicates. Each individual replicate consists of 4,000 genes, 3,600 cells, separated into 3 clusters, and two groups of 3 samples each, corresponding to an average of 200 cells per sample in each cluster. Circles indicate observed FDR for 0.01, 0.05, 0.1 and 0.2 significance thresholds. Note that *scDD* methods were excluded from this analysis because *scDD* does not directly handle covariates.

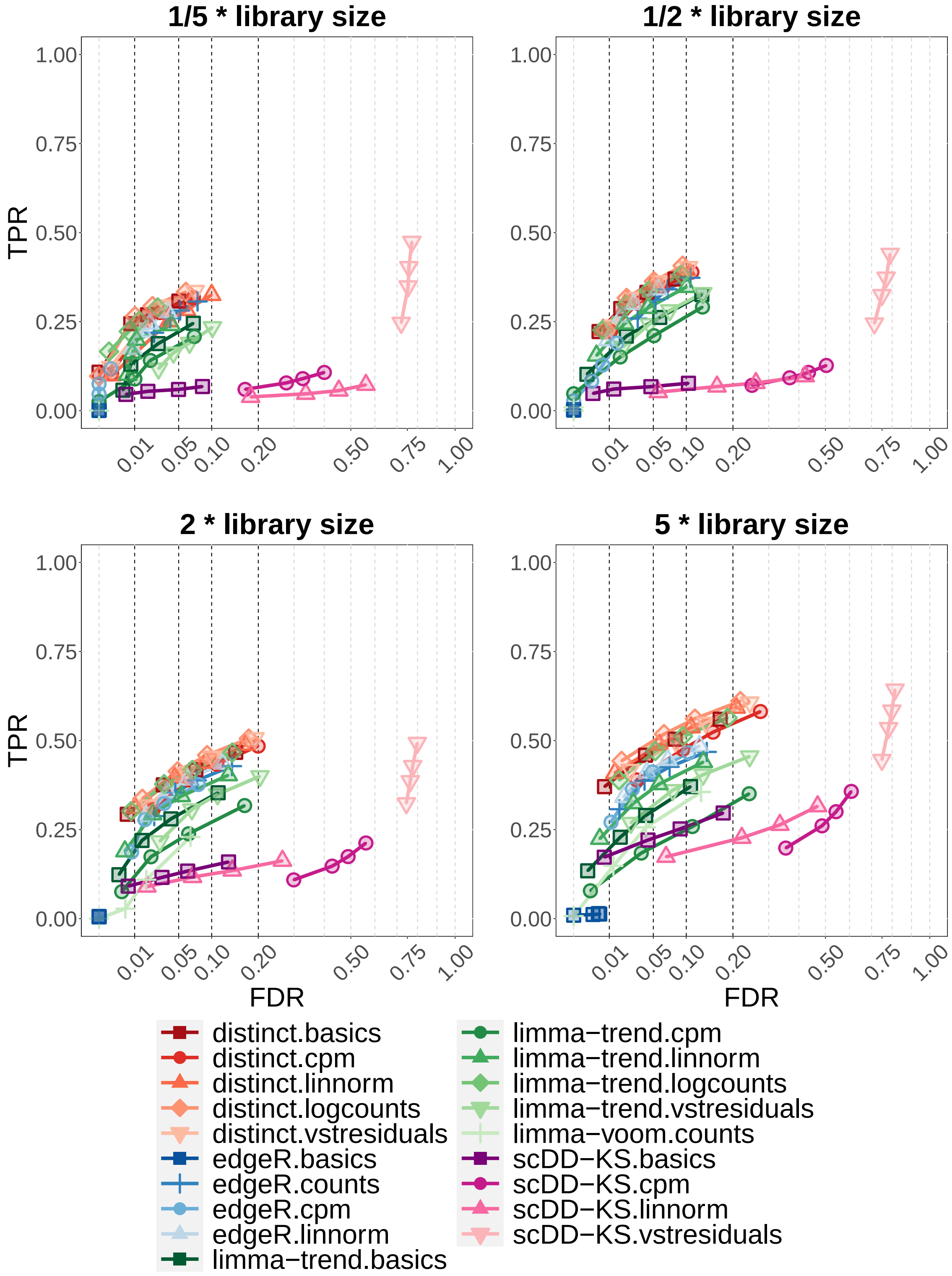


**Supplementary Figure 4:** TPR *vs.* FDR in *muscat* simulated data; with library size 1/5, 1/2, 2 and 5 times as big as the one used in the original simulation study (Figure 2 of the main paper). Results are aggregated over the five differential type (DE, DP, DM, DB and DV), contributing in equal fraction with five replicate simulations. Each individual replicate consists of 4,000 genes, 3,600 cells, separated into 3 clusters, and two groups of 3 samples each, corresponding to an average of 200 cells per sample in each cluster. Circles indicate observed FDR for 0.01, 0.05, 0.1 and 0.2 significance thresholds. Note that *scDD-perm* and MM were excluded from this analysis due to their computational cost.

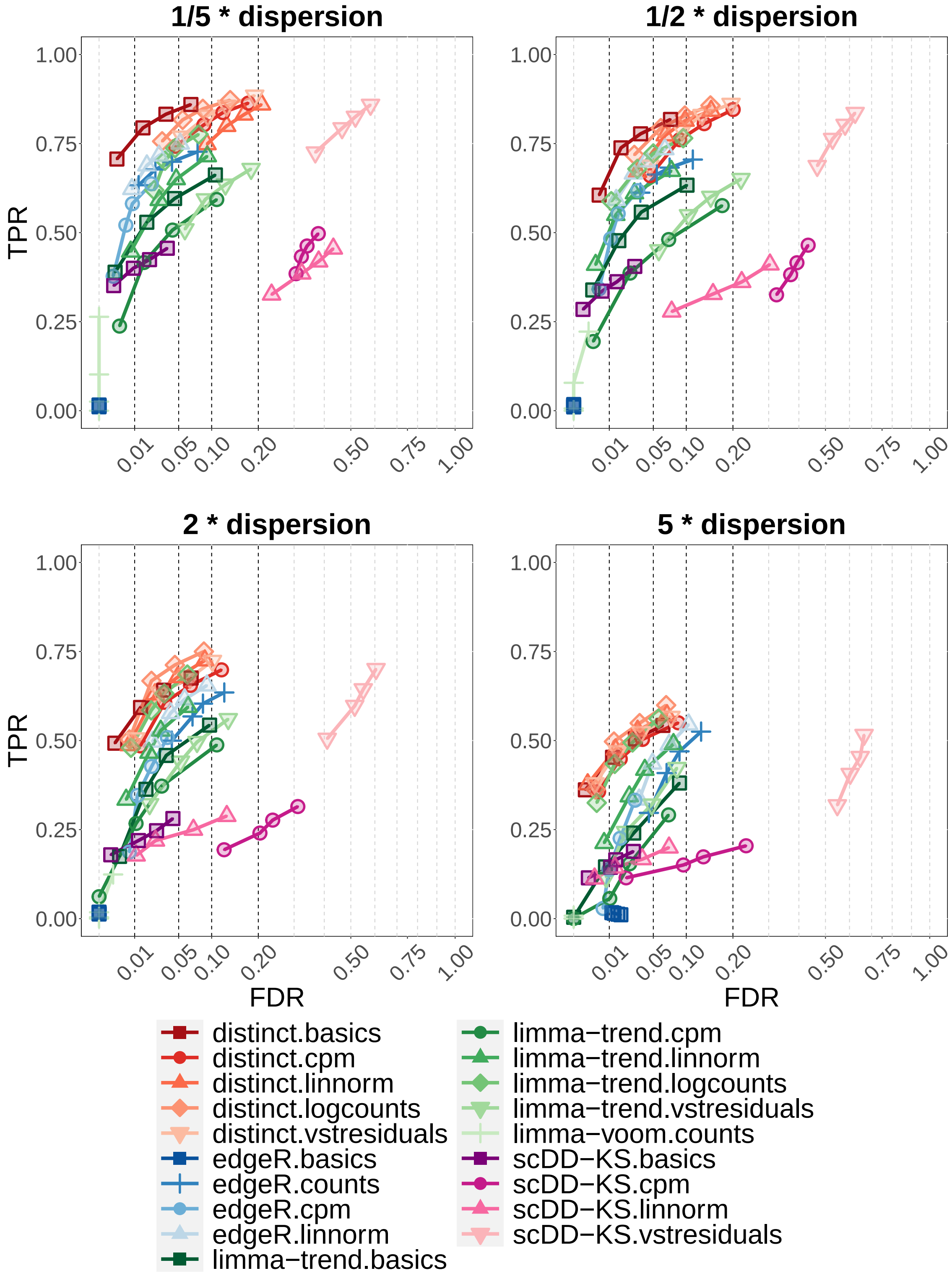


**Supplementary Figure 5:** TPR *vs.* FDR in *muscat* simulated data; with dispersion parameter 1/5, 1/2, 2 and 5 times as big as the one used in the original simulation study (Figure 2 of the main paper). Results are aggregated over the five differential type (DE, DP, DM, DB and DV), contributing in equal fraction with five replicate simulations. Each individual replicate consists of 4,000 genes, 3,600 cells, separated into 3 clusters, and two groups of 3 samples each, corresponding to an average of 200 cells per sample in each cluster. Circles indicate observed FDR for 0.01, 0.05, 0.1 and 0.2 significance thresholds. Note that *scDD-perm* and MM were excluded from this analysis due to their computational cost.

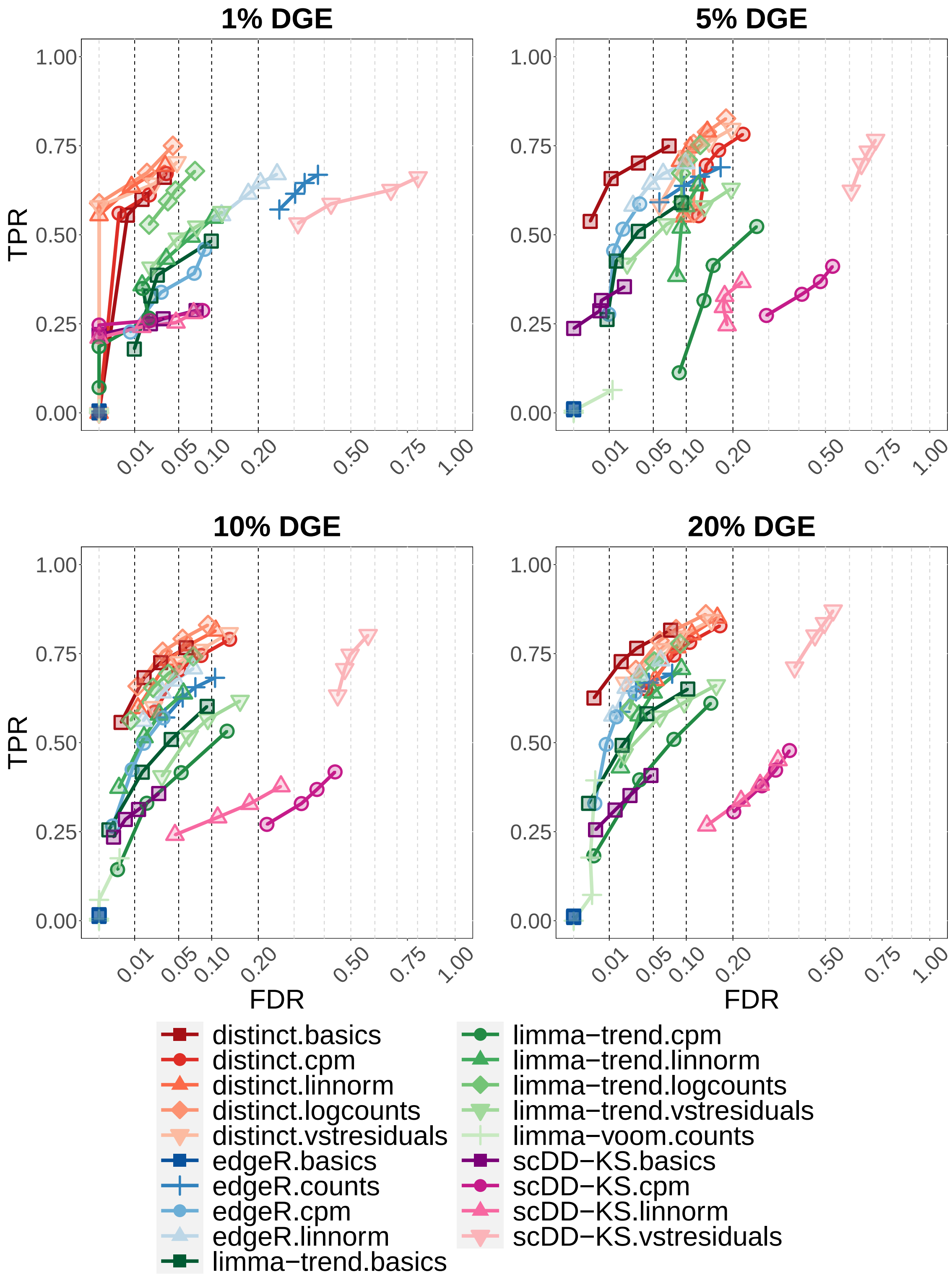


**Supplementary Figure 6:** TPR *vs.* FDR in *muscat* simulated data; with 1%, 5%, 10% (as in the original simulation study in Figure 2 of the main paper), and 20% of significant genes. Results are aggregated over the five differential type (DE, DP, DM, DB and DV), contributing in equal fraction with five replicate simulations. Each individual replicate consists of 4,000 genes, 3,600 cells, separated into 3 clusters, and two groups of 3 samples each, corresponding to an average of 200 cells per sample in each cluster. Circles indicate observed FDR for 0.01, 0.05, 0.1 and 0.2 significance thresholds. Note that *scDD-perm* and MM were excluded from this analysis due to their computational cost.

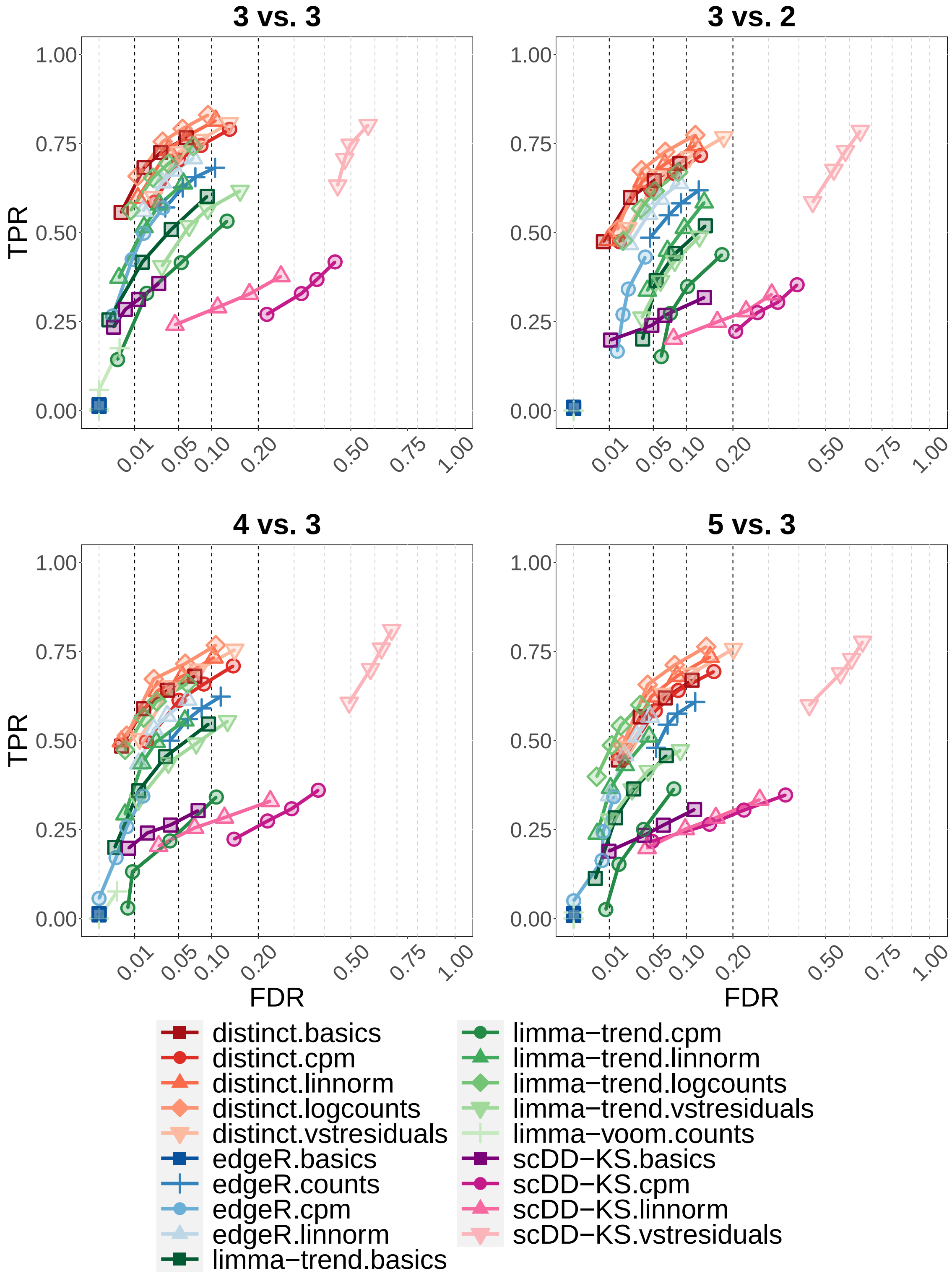


**Supplementary Figure 7:** TPR *vs.* FDR in *muscat* simulated data, considered various unbalanced designs by comparing two groups of different sample sizes: 3 *vs.* 3 (as in the original simulation study in Figure 2 of the main paper), 3 *vs.* 2, 4 *vs.* 3, and 5 *vs.* 3. Results are aggregated over the five differential type (DE, DP, DM, DB and DV), contributing in equal fraction with five replicate simulations. Each individual replicate consists of 4,000 genes, 3,600 cells, separated into 3 clusters, and two groups of 3 samples each, corresponding to an average of 200 cells per sample in each cluster. Circles indicate observed FDR for 0.01, 0.05, 0.1 and 0.2 significance thresholds. Note that *scDD-perm* and MM were excluded from this analysis due to their computational cost.

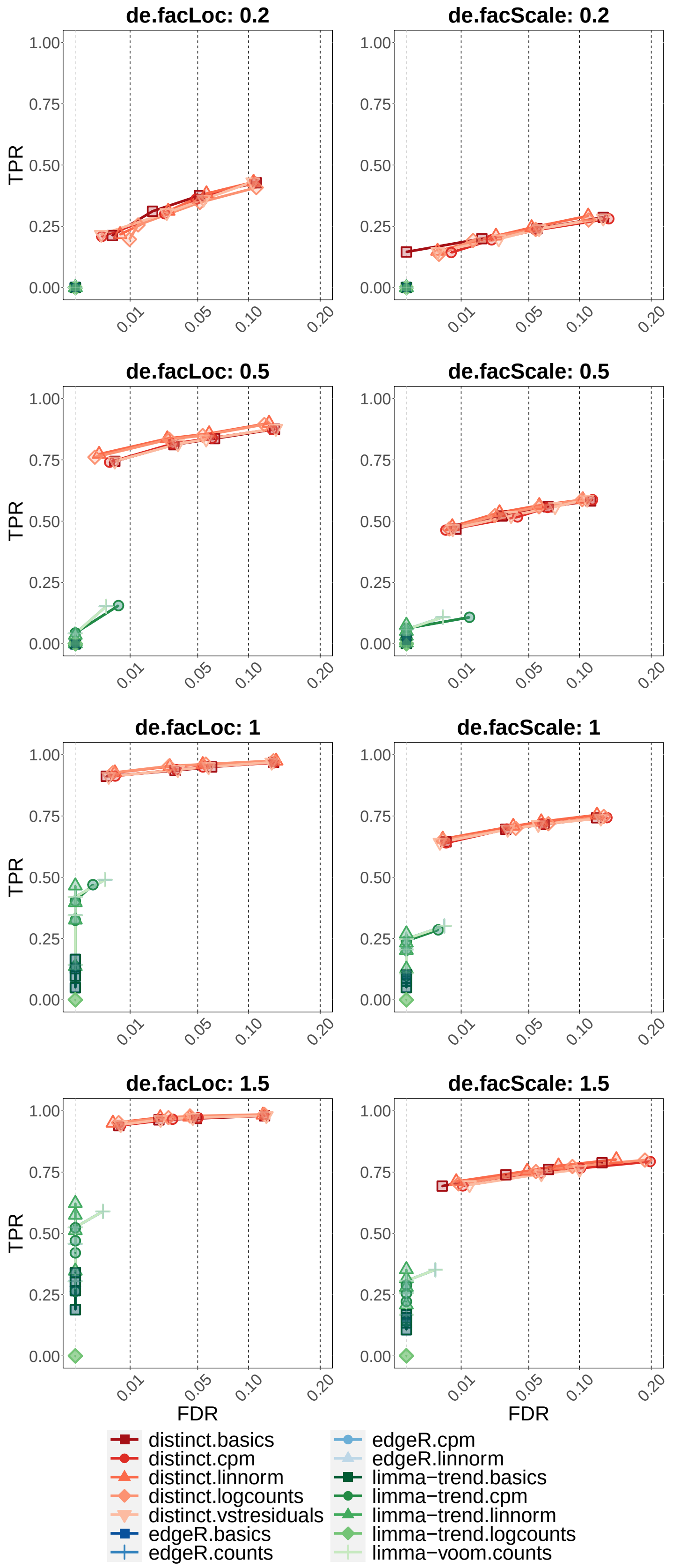


**Supplementary Figure 8:** TPR *vs.* FDR in *SplatPop* simulated data with batch effects, with various degrees of differential location (left) and scale (right) simulations, primarily affecting the mean and variance, respectively. The differential scale and location parameters between batches and groups were identical (i.e., increasing together from 0.2 to 1.5). Circles indicate observed FDR for 0.01, 0.05, 0.1 and 0.2 significance thresholds. Each simulation consists of 20,345 genes genes, 800 cells (belonging to the same cluster), and two groups of 4 samples each, corresponding to an average of 100 cells per sample.

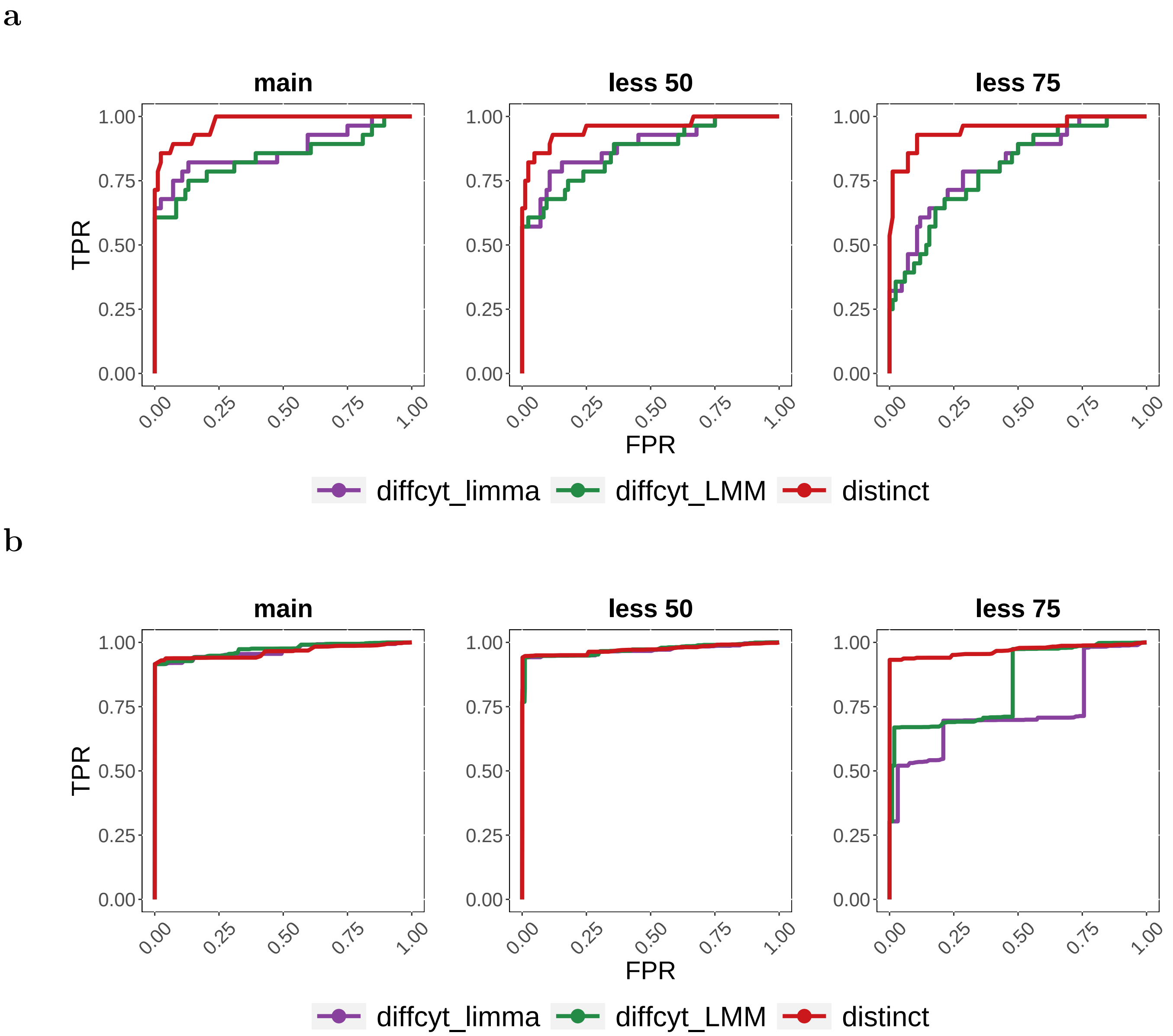


**Supplementary Figure 9:** Receiver operating characteristic (ROC) curve in *diffcyt* non-null semi-simulated data. Each simulation consists of 88,435 cells and two groups of 8 samples each. 'main', 'less 50' and 'less 75' indicate the main simulation, and those where differential effects are diluted by 50 and 75%, respectively. We evaluated methods' performance in terms of detecting DS for phosphorylated ribosomal protein S6 (pS6) in B cells, which is the strongest differential signal across the cell types in this dataset [5, 6]. **(a)** As in the *muscat* simulation study, cells were clustered based on manually annotated cell types [6]. **(b)** As in Weber *et al.* [6], cells were clustered in an unsupervised manner.

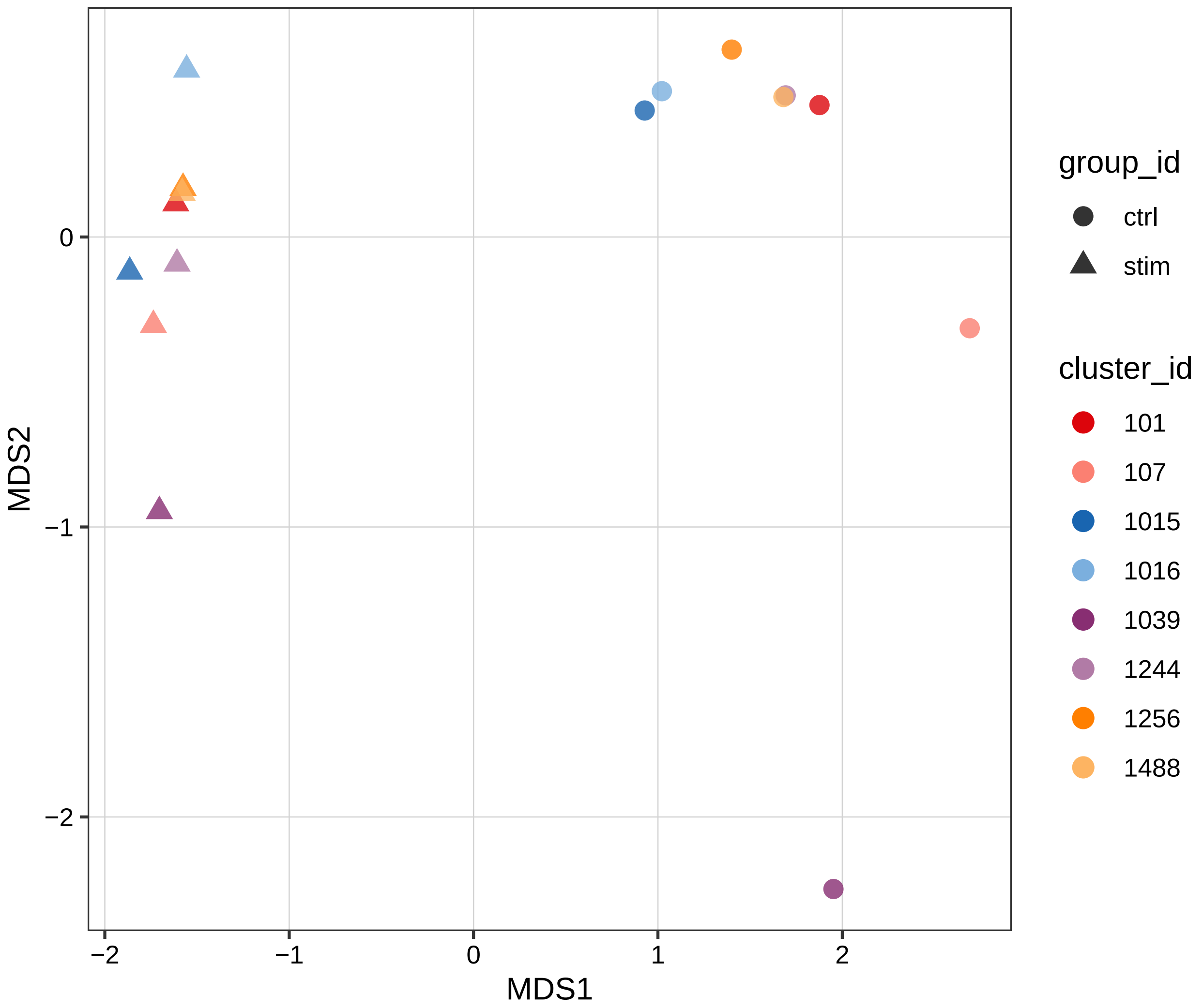


**Supplementary Figure 10:** Multidimensional scaling (MDS) plot, performed via *muscat*'s *pbMDS* function [1], of the pseudo-bulk counts per million (CPM) from the *Kang* dataset [3]. Legend label *cluster_ id* refers to the id of samples, where sample 1039 emerges as a potential outlier in both control and stimulated conditions.

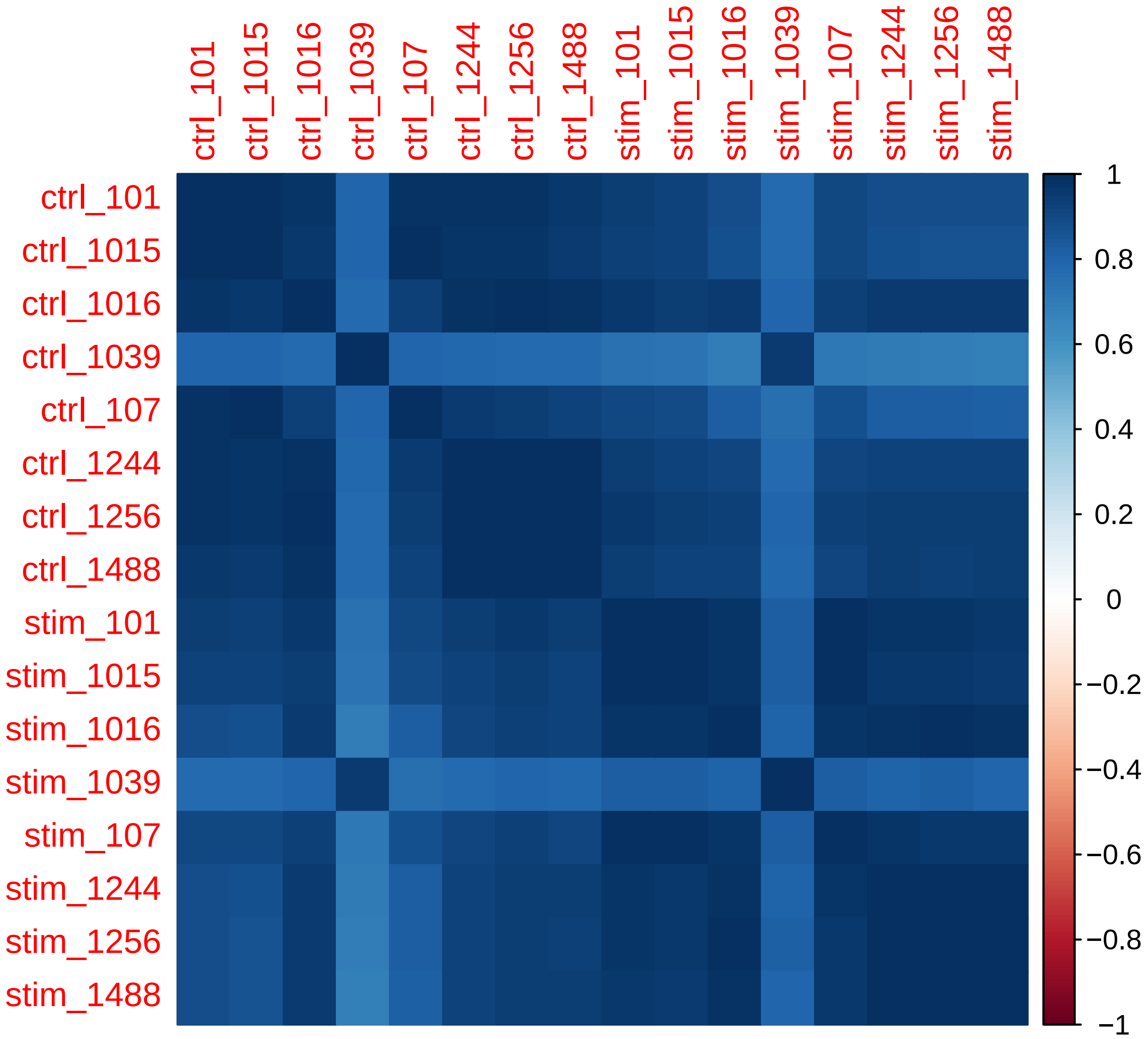


**Supplementary Figure 11:** Visual representation of the correlation, between samples, of the pseudo-bulk CPMs from the *Kang* dataset [3]. Again, sample 1039 behaves as a potential outlier in both control (ctrl_1039) and stimulated (stim_1039) conditions.

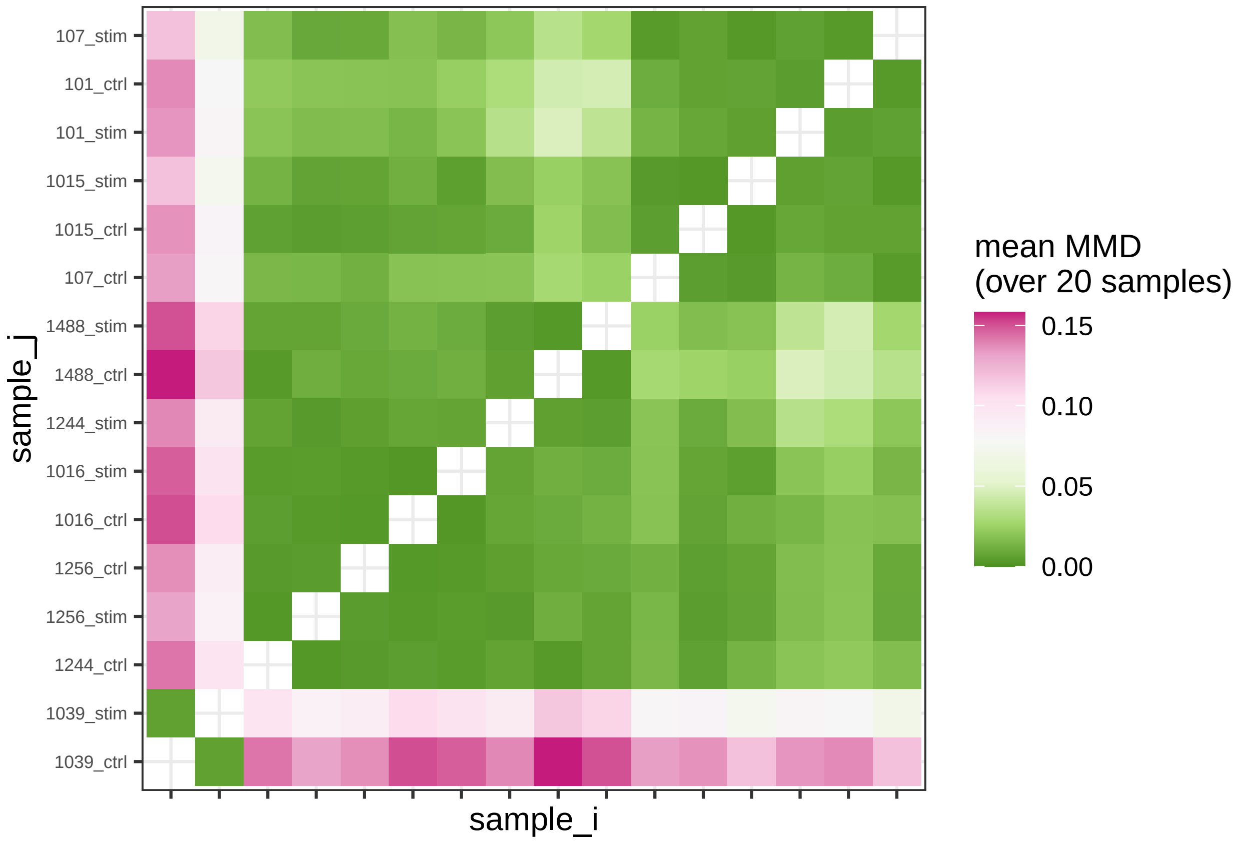


**Supplementary Figure 12:** Maximum mean discrepancy (MMD) of quality controls of samples, performed via *SampleQC*'s *plot_mmd_heatmap* function [4], from the *Kang* dataset [3]. Again, sample 1039 appears to be a potential outlier in both control (ctrl_1039) and stimulated (stim_1039) conditions.

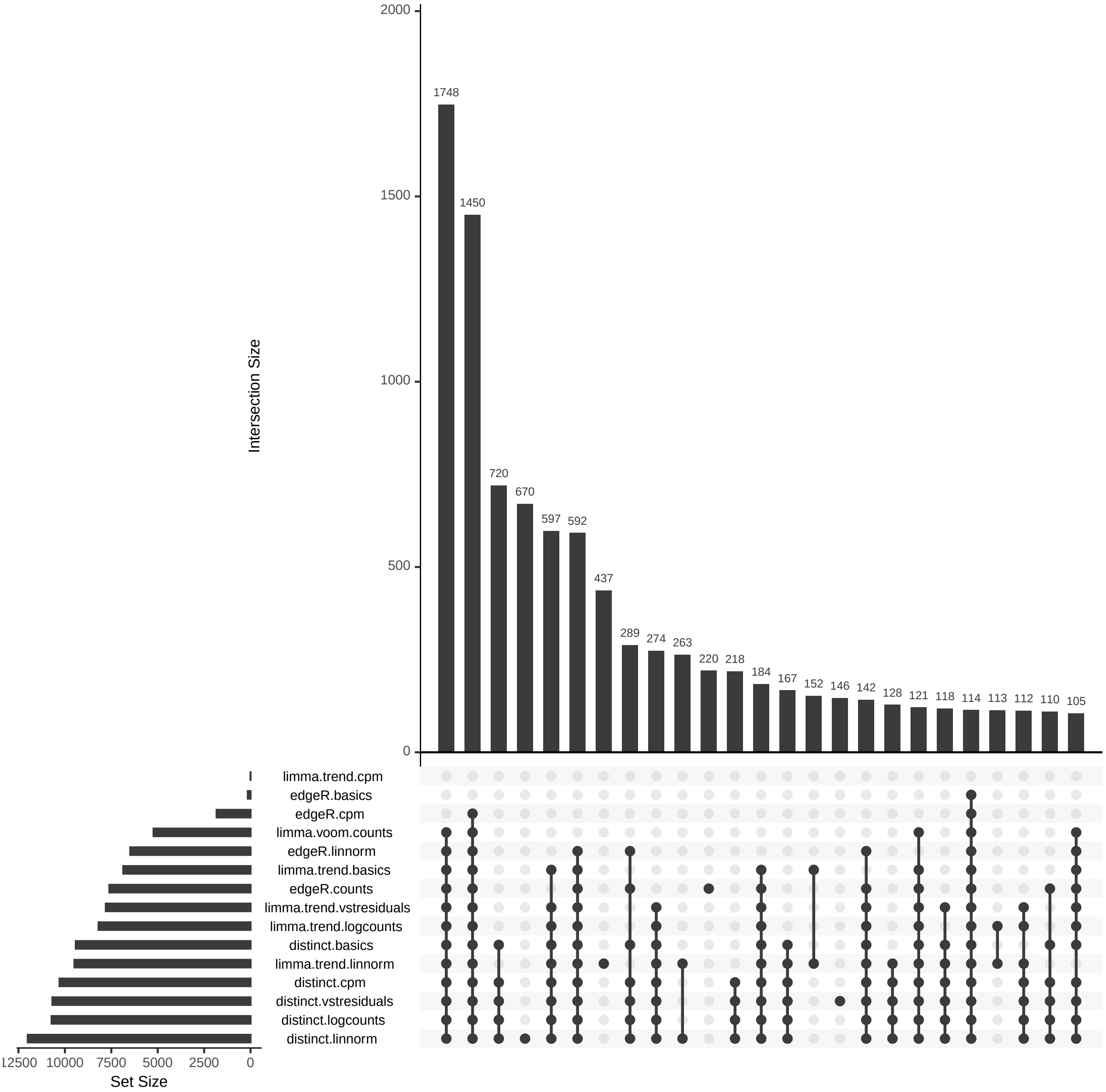


**Supplementary Figure 13:** Visualization (obtained via *UpSetR* R package [2]) of set intersections between differential patterns identified by each method on the *Kang* dataset, when comparing controls and stimulated samples (adjusted p-value $< 0.05$). *distinct* identifies more patterns than PB methods, with *edgeR* based on CPMs being the most conservative approach. Note that *distinct* results are highly coherent across different input data.

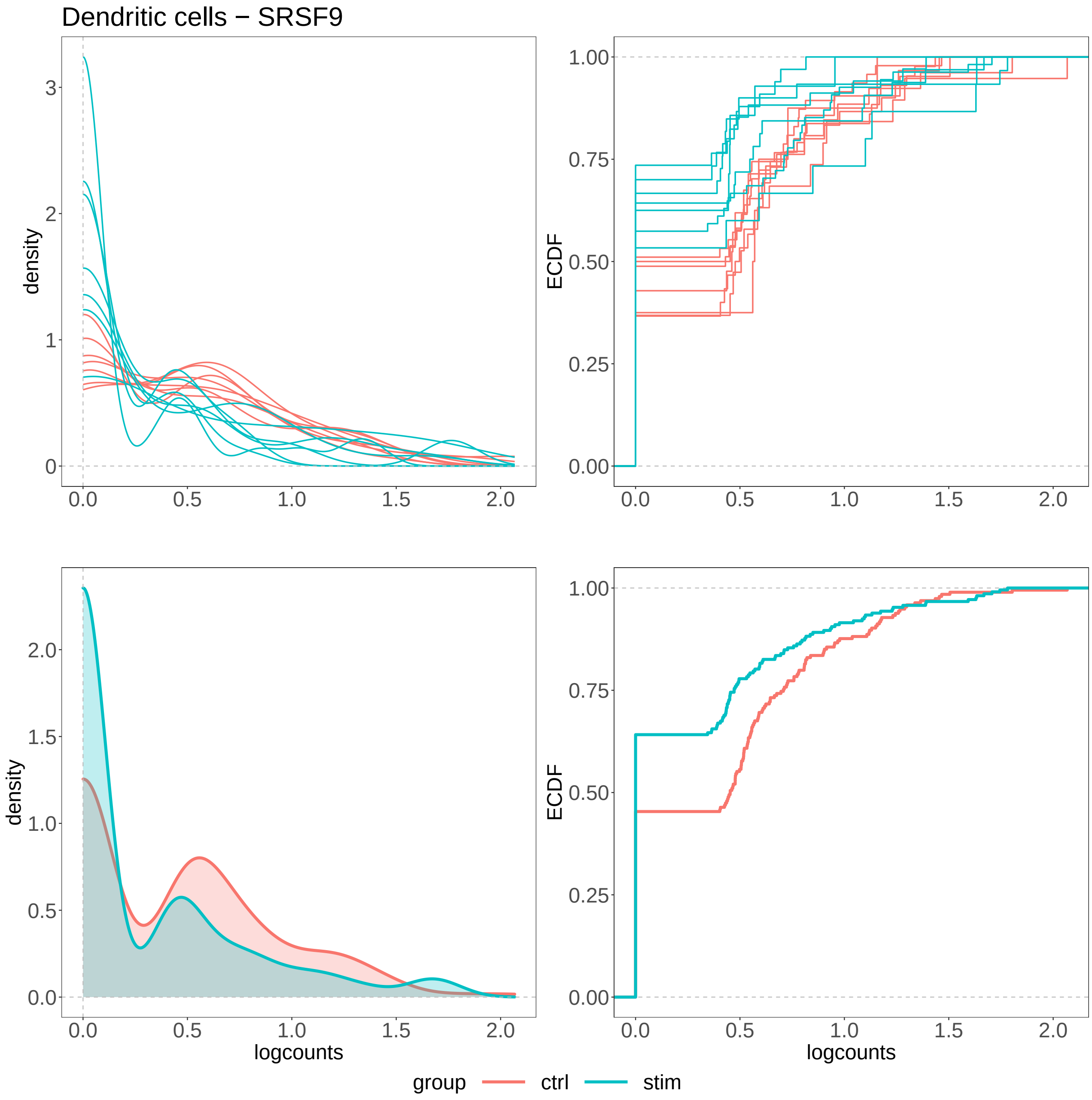


**Supplementary Figure 14:** Density (left panels) and ECDF (right panels), for the logcounts of gene SRSF9 in Dendritic cells, resembling a DP differential pattern. Top panels display one line per sample, while bottom panels show results aggregated at the group level. This result is identified by *distinct* on all 5 input data (adjusted p-value $< 0.1$), and not by any PB tool (adjusted p-value $> 0.1$), on the *Kang* dataset, when comparing controls and stimulated samples.

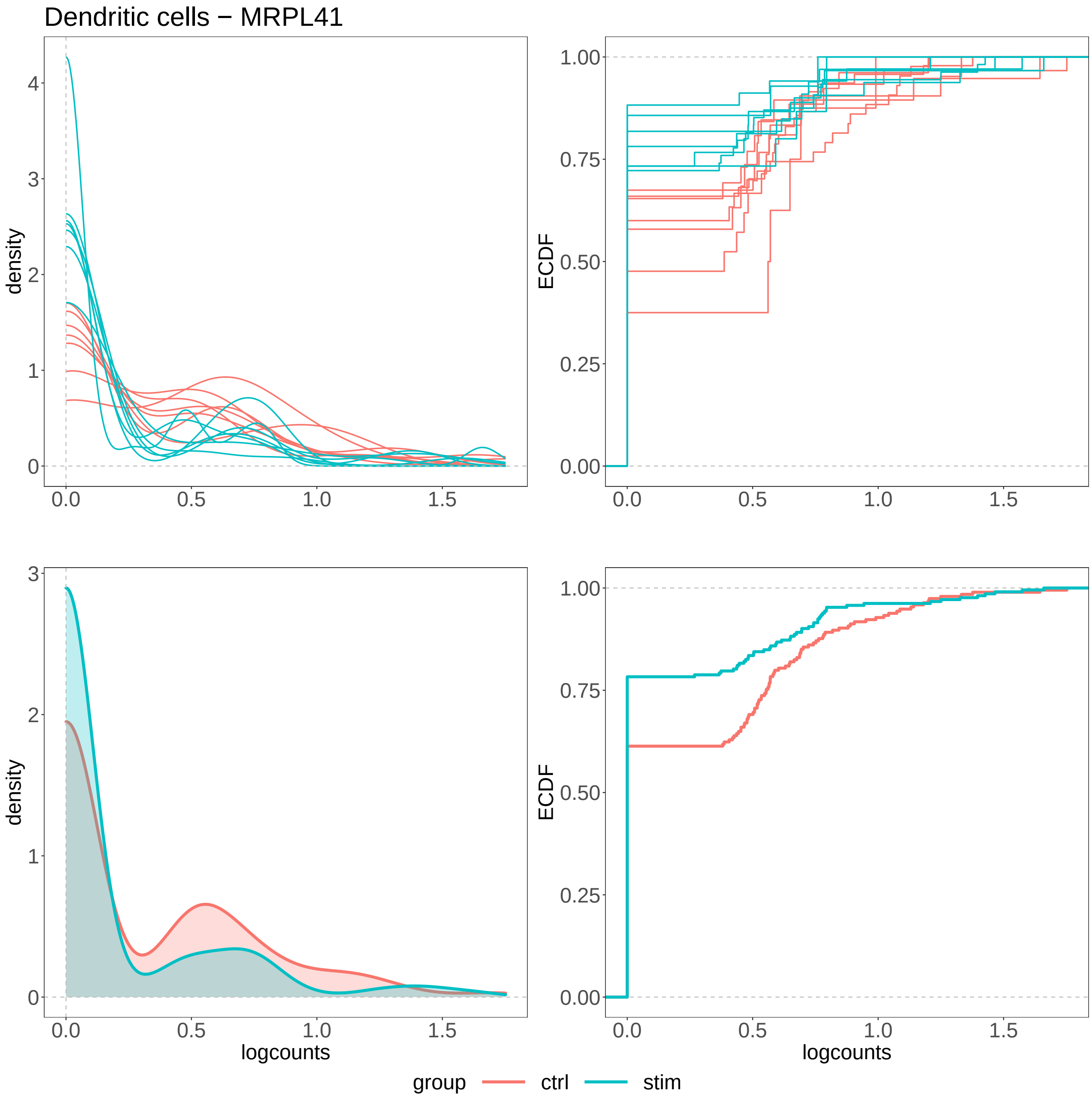


**Supplementary Figure 15:** Density (left panels) and ECDF (right panels), for the logcounts of gene MRPL41 in Dendritic cells, resembling a DP differential pattern. Top panels display one line per sample, while bottom panels show results aggregated at the group level. This result is identified by *distinct* on all 5 input data (adjusted p-value $< 0.1$), and not by any PB tool (adjusted p-value $> 0.1$), on the *Kang* dataset, when comparing controls and stimulated samples.

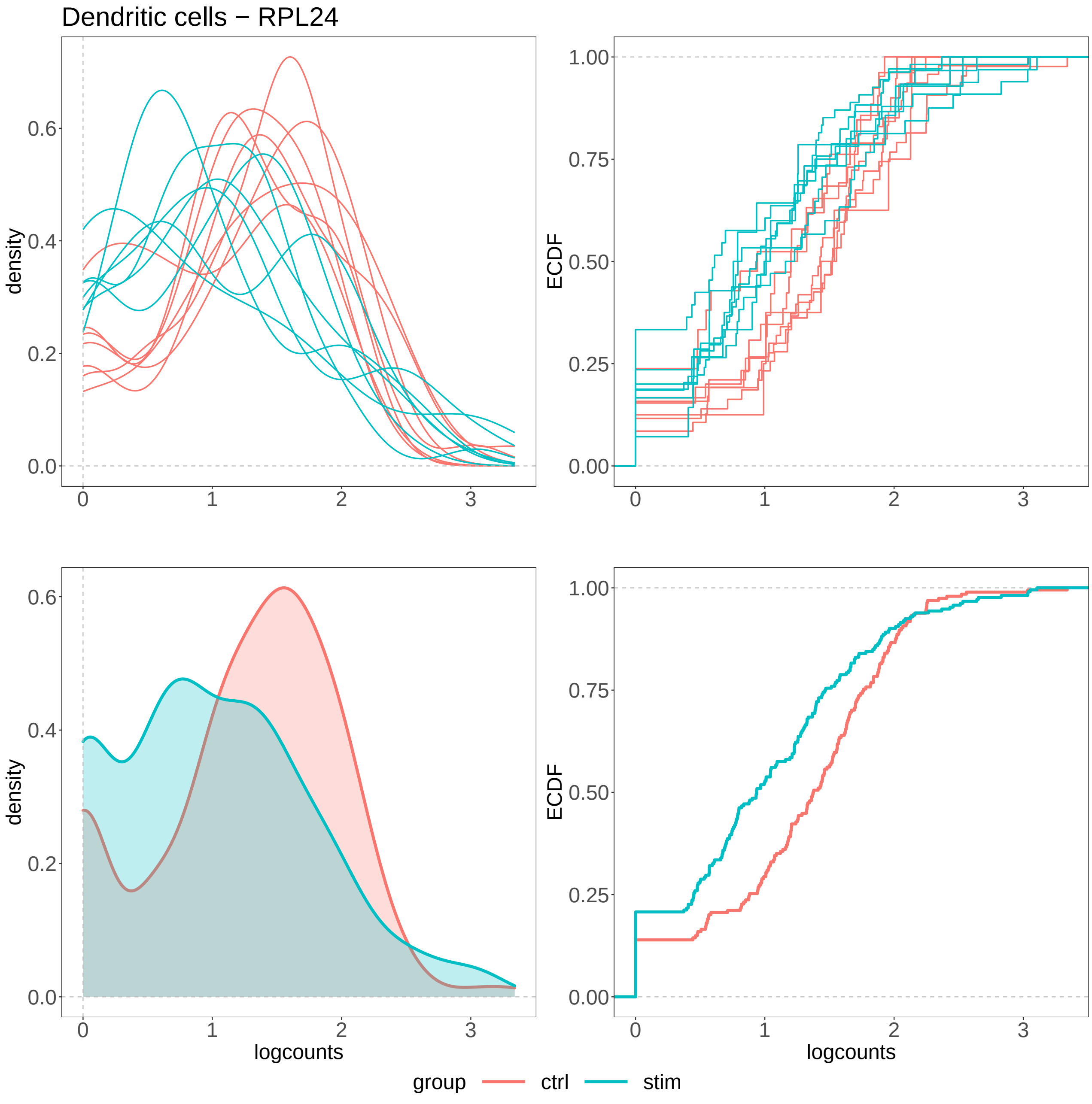


**Supplementary Figure 16:** Density (left panels) and ECDF (right panels), for the logcounts of gene RPL24 in Dendritic cells, resembling a DP differential pattern. Top panels display one line per sample, while bottom panels show results aggregated at the group level. This result is identified by *distinct* on all 5 input data (adjusted p-value $< 0.1$), and not by any PB tool (adjusted p-value $> 0.1$), on the *Kang* dataset, when comparing controls and stimulated samples.

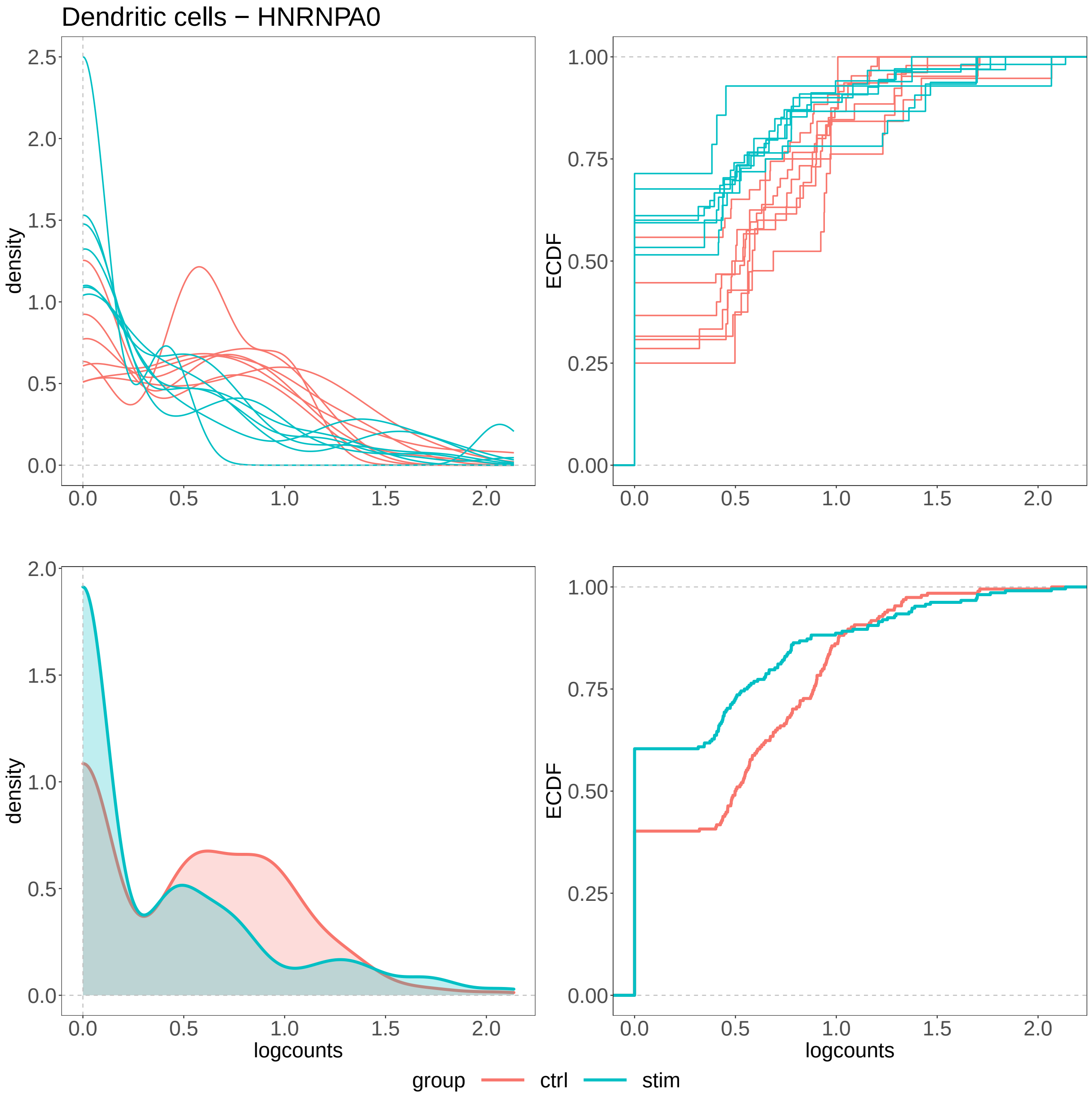


**Supplementary Figure 17:** Density (left panels) and ECDF (right panels), for the logcounts of gene HNRNPA0 in Dendritic cells, resembling a DP differential pattern. Top panels display one line per sample, while bottom panels show results aggregated at the group level. This result is identified by *distinct* on all 5 input data (adjusted p-value $< 0.1$), and not by any PB tool (adjusted p-value $> 0.1$), on the *Kang* dataset, when comparing controls and stimulated samples.

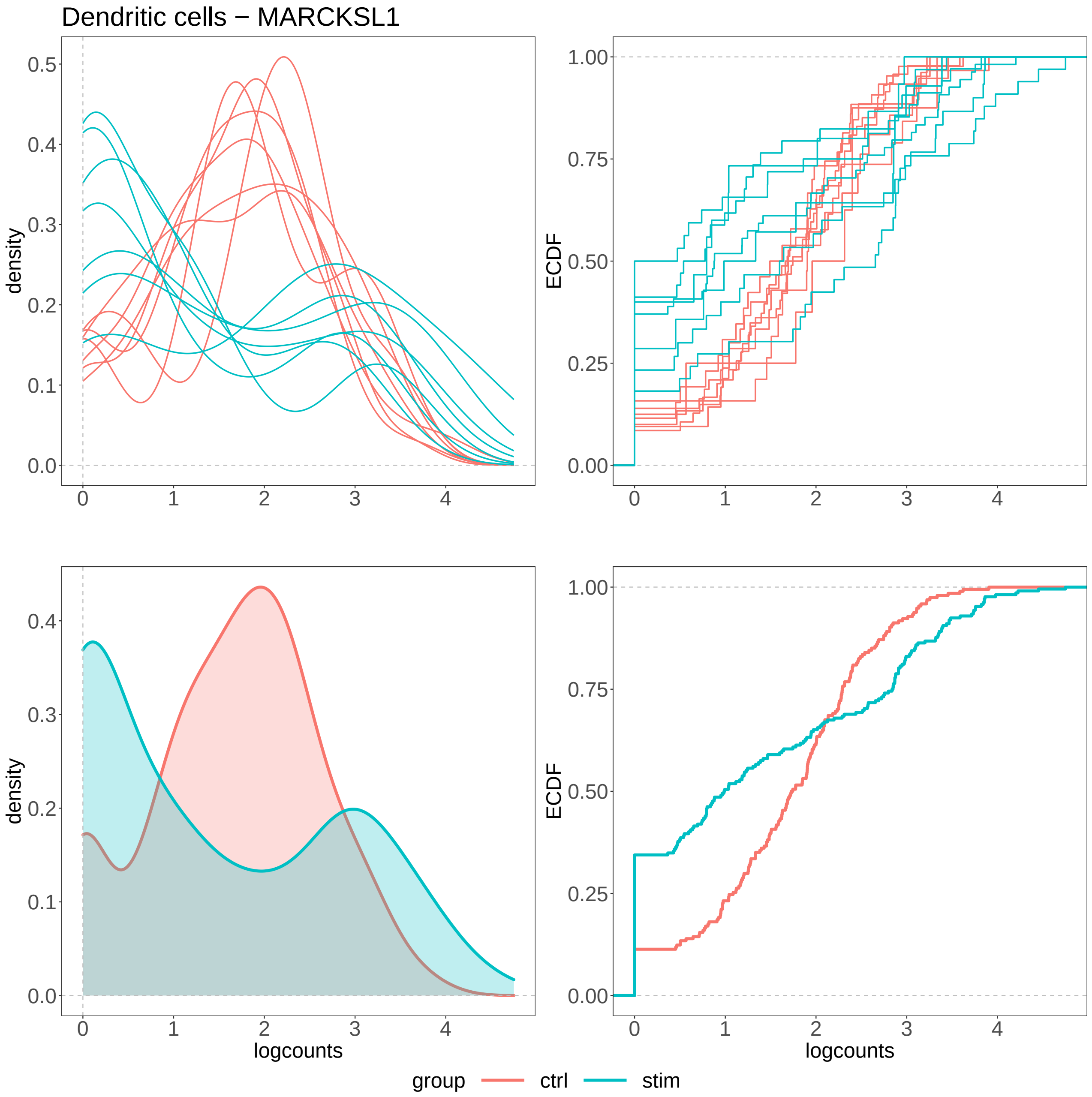


**Supplementary Figure 18:** Density (left panels) and ECDF (right panels), for the logcounts of gene MARCKSL1 in Dendritic cells, resembling a DB differential pattern. Top panels display one line per sample, while bottom panels show results aggregated at the group level. This result is identified by *distinct* on all 5 input data (adjusted p-value $< 0.1$), and not by any PB tool (adjusted p-value $> 0.1$), on the *Kang* dataset, when comparing controls and stimulated samples.

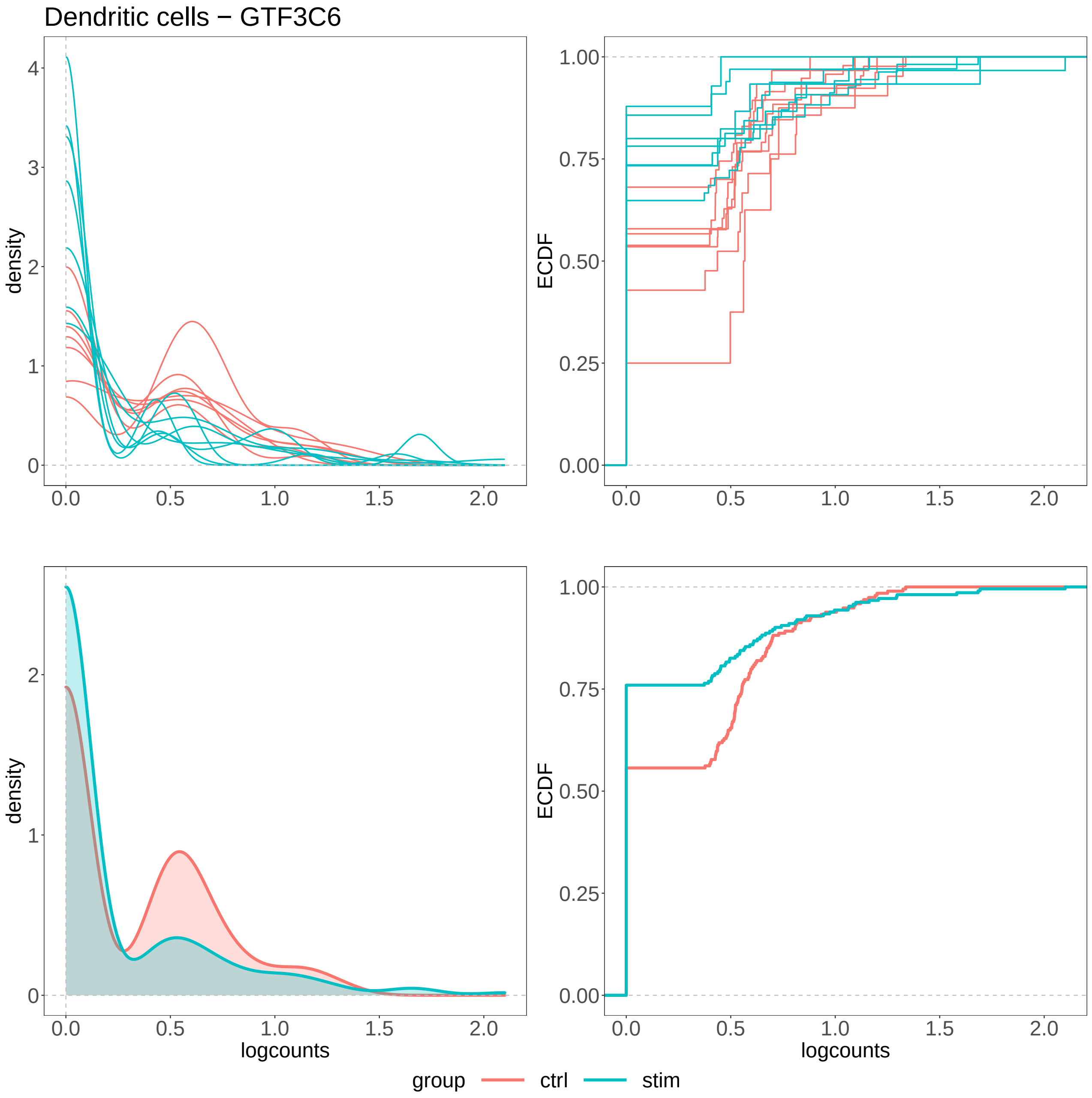


**Supplementary Figure 19:** Density (left panels) and ECDF (right panels), for the logcounts of gene GTF3C6 in Dendritic cells, resembling a DP differential pattern. Top panels display one line per sample, while bottom panels show results aggregated at the group level. This result is identified by *distinct* on all 5 input data (adjusted p-value $< 0.1$), and not by any PB tool (adjusted p-value $> 0.1$), on the *Kang* dataset, when comparing controls and stimulated samples.

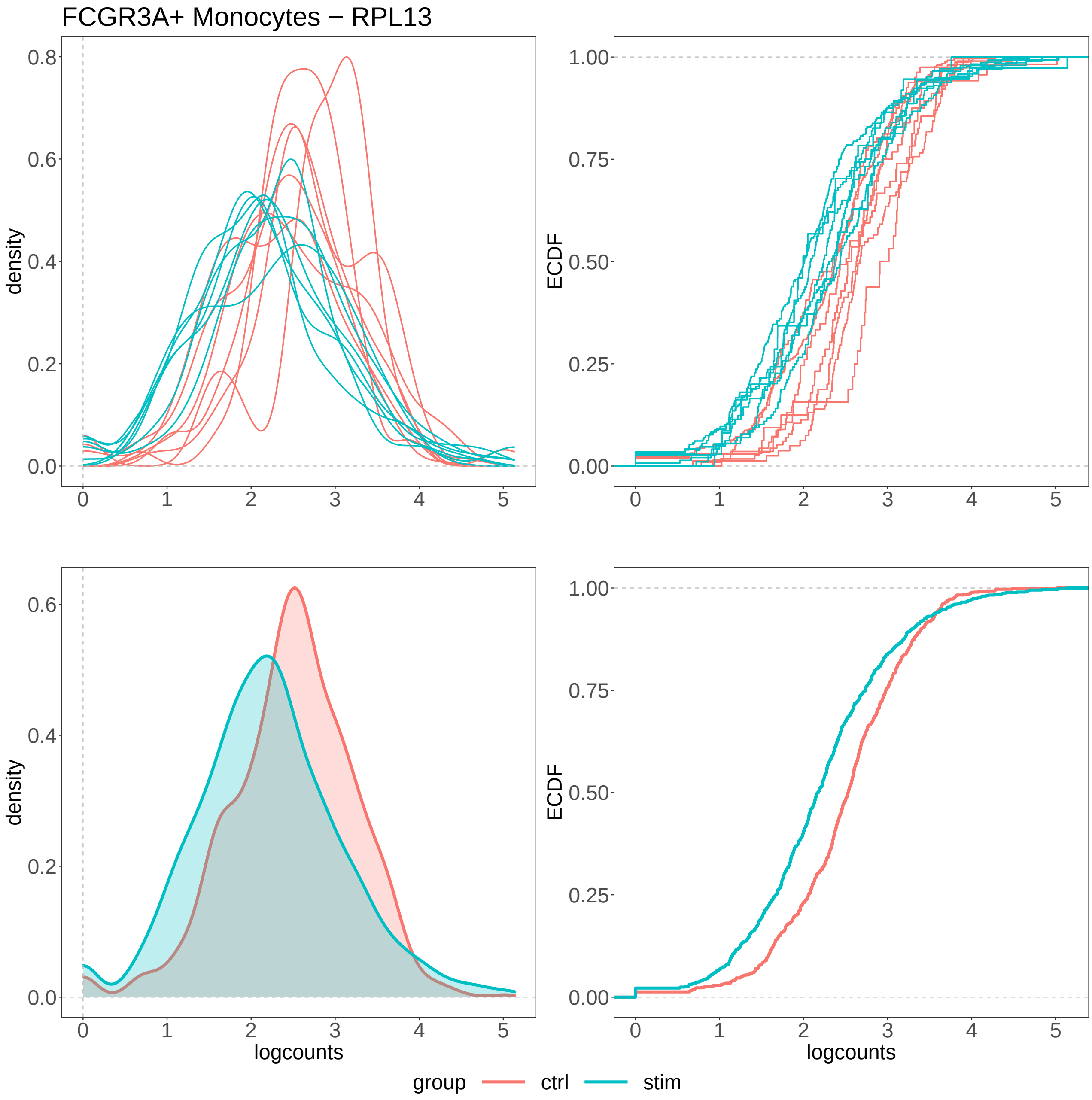


**Supplementary Figure 20:** Density (left panels) and ECDF (right panels), for the logcounts of gene RPL13 in FCGR3A+ Monocytes cells, resembling a DE differential pattern. Top panels display one line per sample, while bottom panels show results aggregated at the group level. This result is identified by *distinct* on all 5 input data (adjusted p-value $< 0.1$), and not by any PB tool (adjusted p-value $> 0.1$), on the *Kang* dataset, when comparing controls and stimulated samples.

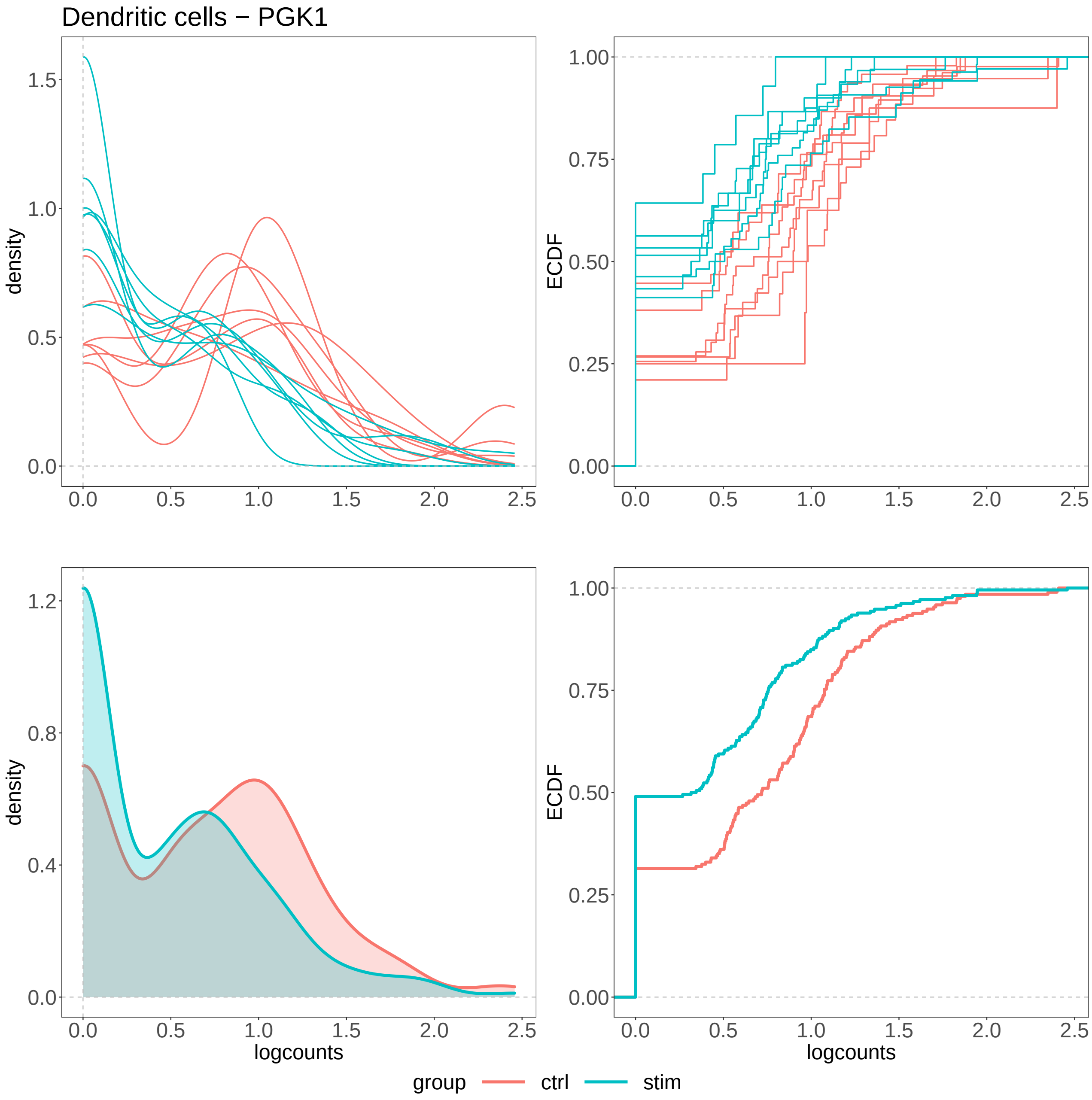


**Supplementary Figure 21:** Density (left panels) and ECDF (right panels), for the logcounts of gene PGK1 in Dendritic cells, resembling a DP differential pattern. Top panels display one line per sample, while bottom panels show results aggregated at the group level. This result is identified by *distinct* on all 5 input data (adjusted p-value $< 0.1$), and not by any PB tool (adjusted p-value $> 0.1$), on the *Kang* dataset, when comparing controls and stimulated samples.

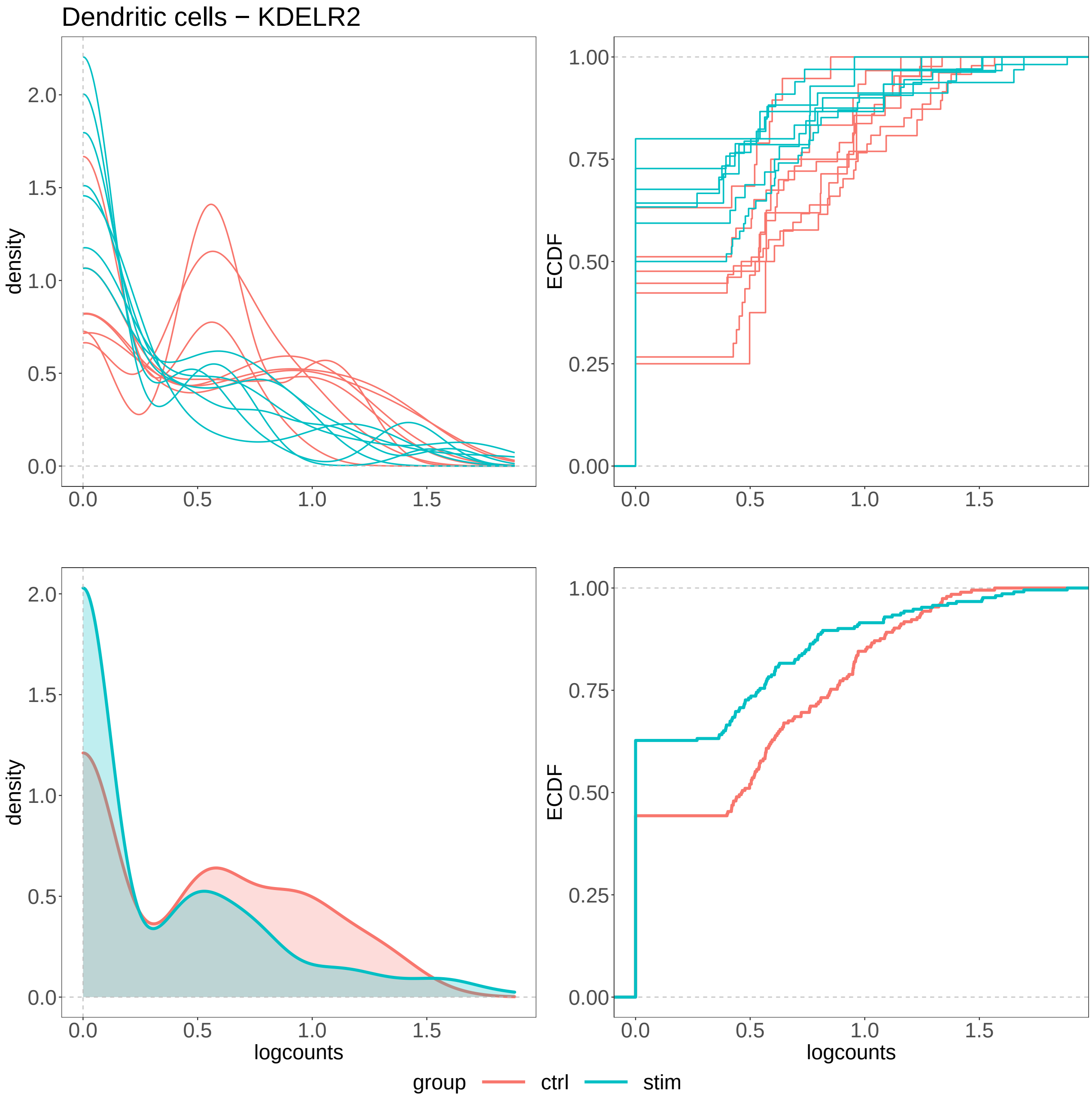


**Supplementary Figure 22:** Density (left panels) and ECDF (right panels), for the logcounts of gene KDELR2 in Dendritic cells, resembling a DP differential pattern. Top panels display one line per sample, while bottom panels show results aggregated at the group level. This result is identified by *distinct* on all 5 input data (adjusted p-value $< 0.1$), and not by any PB tool (adjusted p-value $> 0.1$), on the *Kang* dataset, when comparing controls and stimulated samples.

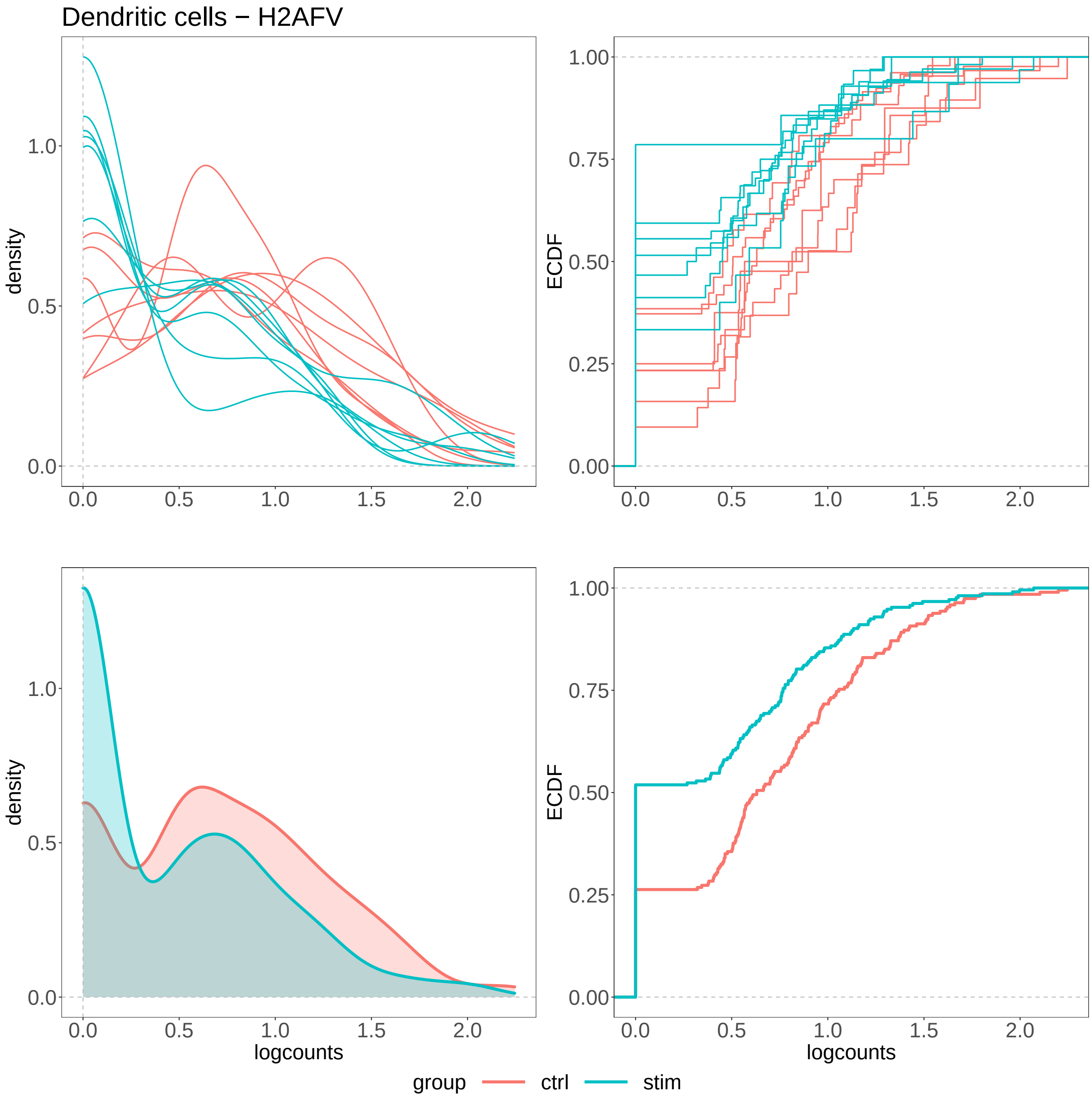


**Supplementary Figure 23:** Density (left panels) and ECDF (right panels), for the logcounts of gene H2AFV in Dendritic cells, resembling a DP differential pattern. Top panels display one line per sample, while bottom panels show results aggregated at the group level. This result is identified by *distinct* on all 5 input data (adjusted p-value $< 0.1$), and not by any PB tool (adjusted p-value $> 0.1$), on the *Kang* dataset, when comparing controls and stimulated samples.

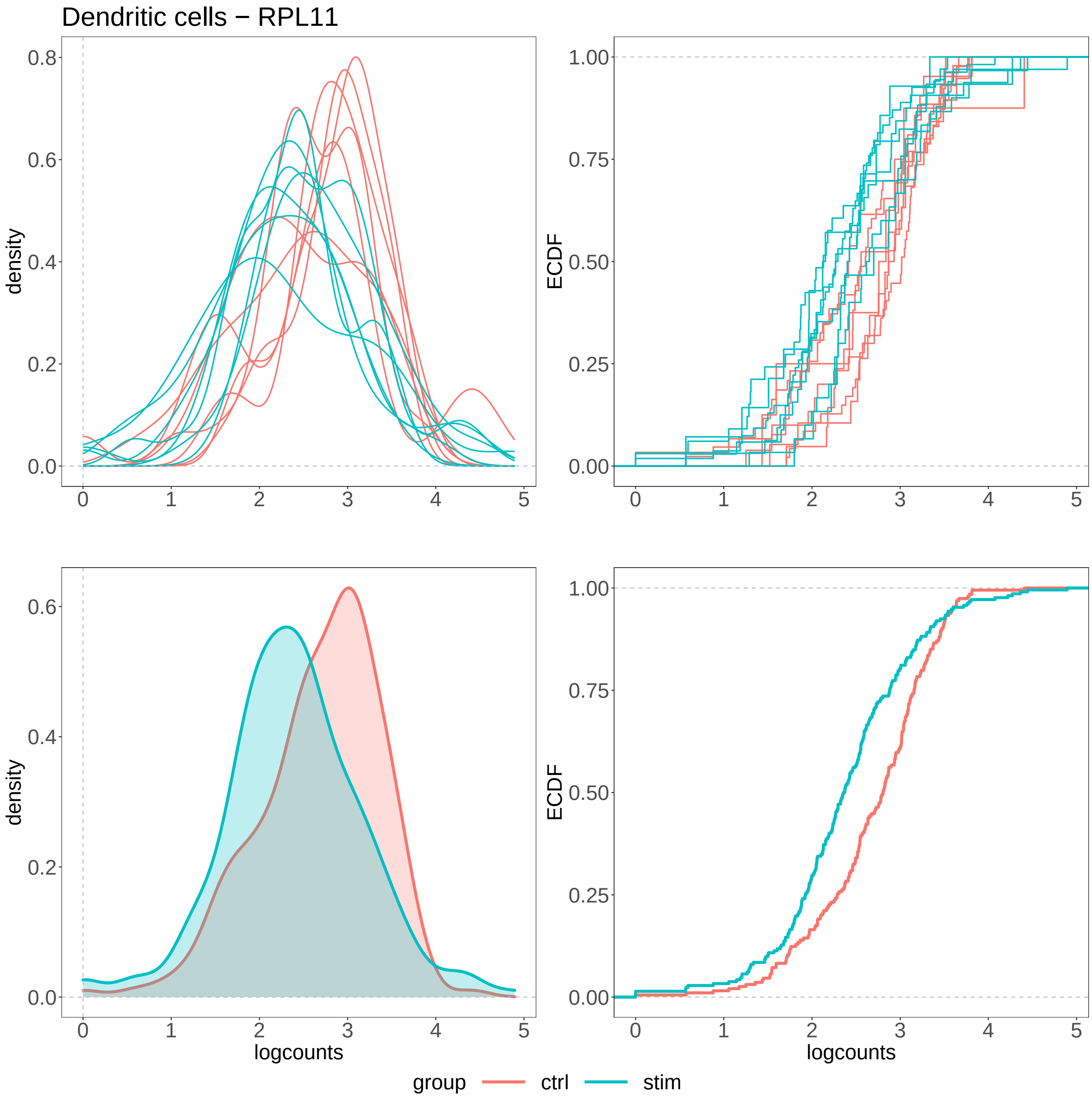


**Supplementary Figure 24:** Density (left panels) and ECDF (right panels), for the logcounts of gene RPL11 in Dendritic cells, resembling a DE differential pattern. Top panels display one line per sample, while bottom panels show results aggregated at the group level. This result is identified by *distinct* on all 5 input data (adjusted p-value $< 0.1$), and not by any PB tool (adjusted p-value $> 0.1$), on the *Kang* dataset, when comparing controls and stimulated samples.

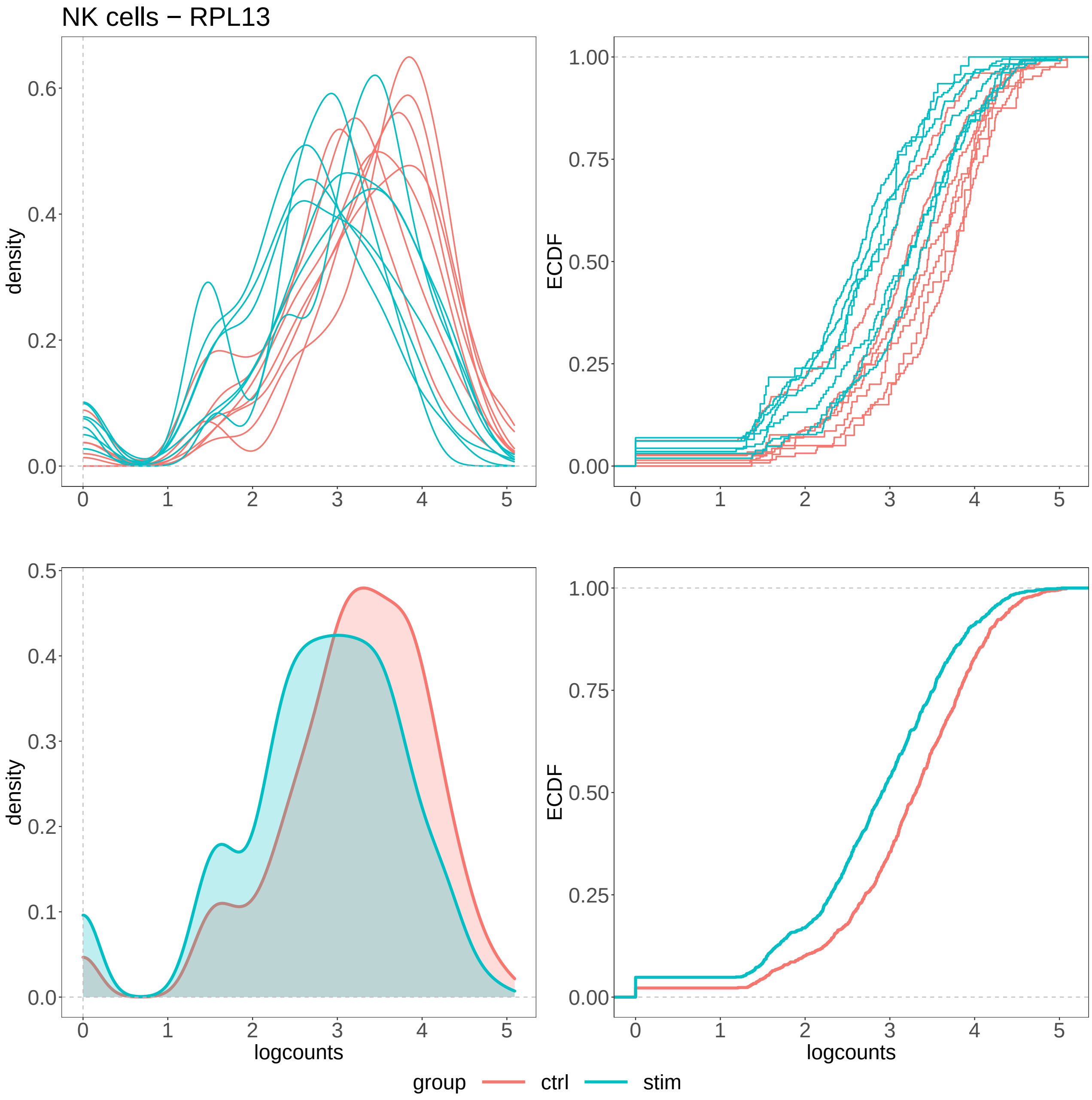


**Supplementary Figure 25:** Density (left panels) and ECDF (right panels), for the logcounts of gene RPL13 in NK cells, resembling a DE differential pattern. Top panels display one line per sample, while bottom panels show results aggregated at the group level. This result is identified by *distinct* on all 5 input data (adjusted p-value $< 0.1$), and not by any PB tool (adjusted p-value $> 0.1$), on the *Kang* dataset, when comparing controls and stimulated samples.
